# Decoding of columnar-level organization across cortical depth using BOLD- and CBV-fMRI at 7 T

**DOI:** 10.1101/2023.09.28.560016

**Authors:** Daniel Haenelt, Denis Chaimow, Marianna Elisa Schmidt, Shahin Nasr, Nikolaus Weiskopf, Robert Trampel

## Abstract

Multivariate pattern analysis (MVPA) methods are a versatile tool to retrieve information from neurophysiological data obtained with functional magnetic resonance imaging (fMRI) techniques. Since fMRI is based on measuring the hemodynamic response following neural activation, the spatial specificity of the fMRI signal is inherently limited by contributions of macrovascular compartments that drain the signal from the actual location of neural activation, making it challenging to image cortical structures at the spatial scale of cortical columns and layers. By relying on information from multiple voxels, MVPA has shown promising results in retrieving information encoded in fine-grained spatial patterns. We examined the spatial specificity of the signal exploited by MVPA. Over multiple sessions, we measured ocular dominance columns (ODCs) in human primary visual cortex (V1) with different acquisition techniques at 7 T. For measurements with blood oxygenation level dependent (BOLD) contrast, we included both gradient echo- (GE-BOLD) and spin echo-based (SE-BOLD) sequences. Furthermore, we acquired data using the vascular-space-occupancy (VASO) fMRI technique, which is sensitive to cerebral blood volume (CBV) changes. We used the data to decode eye-of-origin from signals across cortical layers. While ocularity information can be decoded with all imaging techniques, laminar profiles reveal that macrovascular contributions affect all acquisition methods, limiting their specificity across cortical depth. Therefore, although MVPA is a promising approach for investigating the mesoscopic circuitry of the human cerebral cortex, careful consideration of macrovascular contributions is needed that render the spatial specificity of the extracted signal.

## Introduction

In the cerebral cortex, neurons tend to cluster into functional units across cortical depth (Mountcastle, 1957; Hubel and Wiesel, 1962), which are usually called cortical columns and often denoted as the fundamental building blocks of the cortex (Mountcastle, 1997); however, see (Horton and Adams, 2005) for an alternative perspective. A prominent example is found in the primary visual cortex (V1). V1 mainly receives thalamocortical projections from the lateral geniculate nucleus (LGN) (Wandell, 1995), which contains monocular neurons that are segregated into eye-specific layers (Andrews, Halpern, and Purves, 1997). The monocular information is preserved when entering V1, and projections from the left and right eye are sent to segregated cortical columns, widely known as ocular dominance columns (ODCs) (Hubel and Wiesel, 1969; Tootell et al., 1988; Dougherty et al., 2019), which form a repeating stripes pattern of alternating eye preference (Adams, Sincich, and Horton, 2007).

Functional magnetic resonance imaging (fMRI) is a versatile neuroimaging technique for non-invasive measuring and mapping of brain activity by assessing the hemodynamic response following neural activation (Buxton, 2013). However, due to the limited spatial resolution, conventional fMRI techniques only allow the detection of relatively large pieces of cortex involved in the execution of a specific task (Glover, 2011). Therefore, ODCs with an approximate column width of around 1 mm in humans (Adams, Sincich, and Horton, 2007) and other cortical columns were out of reach for usual fMRI applications.

With the development of MR scanners with higher magnetic field strengths and more sophisticated radiofrequency (RF) coils providing higher signal-to-noise ratio (SNR), mesoscopic structures like ODCs became accessible in humans at the expense of prolonged acquisition times and usage of anisotropic voxels (Menon et al., 1997; Menon and Goodyear, 1999; Dechent and Frahm, 2000; Goodyear and Menon, 2001; Cheng, Waggoner, and Tanaka, 2001; Yacoub et al., 2007). Only with the emergence of ultra-high field MRI at a field strength of 7 Tesla and above, it became possible to measure ODCs with isotropic voxels at sub-millimeter resolution (Nasr, Polimeni, and Tootell, 2016; Feinberg, Vu, and Beckett, 2018; Zaretskaya et al., 2020; Hollander et al., 2021; Akbari et al., 2023; Nasr et al., 2025).

Given the average cortical thickness of 2–4 mm (Fischl and Dale, 2000) and its convoluted structure, the use of isotropic voxels at sub-millimeter resolution is necessary for the reliable sampling of data at different cortical depths (Turner and Geyer, 2014). This recent possibility is intriguing since the cerebral cortex is known to be composed of several layers, e.g., in terms of cytoarchitecture (Brodmann, 1909), myeloarchitecture (Vogt and Vogt, 1919), and vasculature (Duvernoy, Delon, and Vannson, 1981). Furthermore, cortical layers generally differ in their connectivity profile within and to other cortical areas, e.g., feedforward and feedback signaling between cortical areas in a hierarchically organized cortical system (Felleman and Van Essen, 1991). Thus, the mapping of cortical columns at different cortical depths with fMRI enables studying the local microcircuitry of the cerebral cortex *in vivo* (Yang et al., 2021).

The monocular feedforward signal from the LGN enters V1 in layer 4C of corresponding ODCs (Kennedy et al., 1976; Tootell et al., 1988). Layer 4C is located directly below layer 4B, which contains the highly myelinated external band of Baillarger, also called stria of Gennari (Trampel, Ott, and Turner, 2011). Typically, layer 4C is further divided into layers 4Cα and 4Cβ, which receive color-selective parvocellular and “color-blind” magnocellular input from corresponding LGN layers, respectively (Nieuwenhuys, Voogd, and Huijzen, 2008). Above and below layer 4C, the signals from the two eyes converge onto single neurons, which lead to a variable degree of ocularity across cortical depth. However, individual neurons of the same column still tend to receive input predominantly from either the left or right eye, respectively (Wandell, 1995). In this regard, V1 is the first main stage of binocular integration, which is important, for example, for the processing of stereopsis (Poggio, 1995).

However, fMRI provides only an indirect measure of neural activity, most commonly relying on the blood oxygenation level-dependent (BOLD) signal acquired with gradient echo-based sequences (GE-BOLD), which are known to be most sensitive to macrovascular compartments of the cerebral cortex (Turner, 2002), specifically draining veins that carry the deoxygenated blood back to the cortical surface (Polimeni et al., 2010a; Markuerkiaga, Barth, and Norris, 2016). This usually leads to a signal accumulation toward the pial surface, limiting the ability to associate the BOLD response with a specific cortical layer. Alternatively, spin echo-based sequences (SE-BOLD) can be used at high magnetic field strengths (Yacoub et al., 2007). SE-BOLD promises a more specific signal due to the refocusing of extravascular signal contributions from around larger veins (Boxerman et al., 1995). This has the advantage of increasing signal weighting to the microvasculature, which is believed to be closer to the actual location of neural activation. Furthermore, recent advances of imaging approaches with contrast weighted by cerebral blood volume (CBV) using vascular-space-occupancy (VASO) fMRI at higher magnetic fields show promising results in terms of increased laminar specificity (Huber et al., 2017; Huber et al., 2021) at the expense of overall sensitivity.

Next to the choice of the proper acquisition technique, multivariate pattern analysis (MVPA) (Haxby, 2012) methods have been shown to retrieve information manifested in spatial patterns of fMRI activity, which promise increased sensitivity compared to univariate methods (Kriegeskorte and Bandettini, 2007; Formisano and Kriegeskorte, 2012; Vizioli et al., 2020), for example, for the dissociation of bottom-up and top-down processing into different cortical layers (Muckli et al., 2015; Kok et al., 2016; Iamshchinina et al., 2021). However, though the presence of pattern information provides strong evidence for neuronal effects, the spatial scale of the exploited information remains unknown (Formisano and Kriegeskorte, 2012). Interestingly, already at a conventional resolution of 3 × 3 × 3 mm^3^ using GE-BOLD at 3 T, decoding of orientation information is possible from responses in V1 (Haynes and Rees, 2005a; Kamitani and Tong, 2005), which is known to be encoded at a much finer spatial scale at the level of cortical columns (Obermayer and Blasdel, 1993). In the same year, the eye-of-origin could also be decoded from V1 voxels based on a binocular rivalry stimulus (Haynes and Rees, 2005b). These studies started a controversy several years ago (Boynton, 2005; Beeck, 2010; Swisher et al., 2010; Gardner, 2010; Shmuel et al., 2010; Kriegeskorte, Cusack, and Bandettini, 2010; Chaimow et al., 2011; Misaki, Luh, and Bandettini, 2013) about the source of the exploited information. Possible mechanisms were suggested like the aliasing of high spatial frequency information encoded above the Nyquist frequency of the MRI sampling process (Boynton, 2005) (but see (Chaimow et al., 2011)), the contributions from random irregularities of the fine-scale columnar pattern, which lead to information at low spatial frequencies (Haynes and Rees, 2005a; Kamitani and Tong, 2005; Kriegeskorte and Bandettini, 2007) or the exploitation of large-scale information that is not related to the fine-scale columnar pattern (Beeck, 2010). Growing evidence showed that functional biases can also be introduced by large vessels (Turner, 2002; Gardner, 2010; Shmuel et al., 2010; Sengupta et al., 2017), which can be conceptualized as a form of lowpass filtering the neural pattern, which results in a coarser spatial venous pattern (Formisano and Kriegeskorte, 2012). Therefore, neural patterns encoded at the columnar/laminar level might be represented at multiple spatial scales in the fMRI signal (Swisher et al., 2010; Sengupta et al., 2017).

To study the microcircuitry of the cerebral cortex, it is of importance to know the source of the decoded information, e.g., by relating the decoded information to specific cortical layers. In this regard, it might be appealing to use fMRI acquisition techniques that are less sensitive to large vessels in combination with MVPA methods to benefit from the increased sensitivity of multivariate methods, while keeping a high spatial specificity of the exploited signal. However, most decoding studies use the GE-BOLD technique, which is known to be inherently limited by macrovascular contributions, reducing the potential benefits.

In our study, we acquired ODC data from five participants using GE-BOLD, SE-BOLD, and VASO in different sessions to study the laminar specificity of the respective acquisition technique in combination with MVPA to decode the signal of the stimulated eye in V1. Functional data were acquired with nominal isotropic voxel size of 0.8 mm allowing data sampling at different cortical depths.

From the perspective of neural processing, we expected highest eye-of-origin discriminability in deeper cortical layers since eye-specific segregation is most preserved in the input layer 4C. However, due to the drainage of deoxygenated blood toward the pial surface, macrovascular contributions to the fMRI signal were expected to bias the discriminability across cortical depth. Therefore, studying decoding performance of a feedforward signal between acquisition techniques across cortical depth enables the analysis of their different sensitivities to draining vein contributions. We believe that this study gives insights into the capabilities and limitations of using multivariate techniques with different fMRI sequences for disentangling information at the level of cortical layers.

## Materials and methods

### Participants

A total of five healthy volunteers participated in this study, of which two were female (age = 28.00 ± 2.61, mean ± standard deviation in years). Written informed consent was obtained from all participants, and the study received ethical approval from the local ethics committee of the University of Leipzig. All participants had normal or corrected-to-normal visual acuity. We performed the Miles Test (Miles, 1929) with each participant to determine eye dominance, which is stated in ***Supplementary Figure 1***–***Supplementary Figure 5*** for single participants.

### General procedure

Each participant underwent multiple scanning sessions on different days using an ultra-high field (7 T) MRI scanner. The first session was used for reference measurements, during which a high-resolution anatomical reference scan and retinotopy data (Sereno et al., 1995; Engel, Glover, and Wandell, 1997) were acquired. In addition, a high-resolution functional time series without task (GE-BOLD) was obtained using the same parameters as in subsequent functional measurements, in order to aid with between-session registration.

The remaining six sessions were exclusively devoted to ODC mapping (2× GE-BOLD, 2× SE-BOLD, 2× VASO). ***Figure 1*** provides an illustration of slab positioning along with representative temporal signal-to-noise ratio (tSNR) maps for all contrasts. A subset of the retinotopy data had previously been utilized in another experiment (Movahedian Attar et al., 2020), but underwent independent processing for this study. All functional measurements were accompanied by associated field map acquisitions, which were not further used in this project.

**Figure 1.**
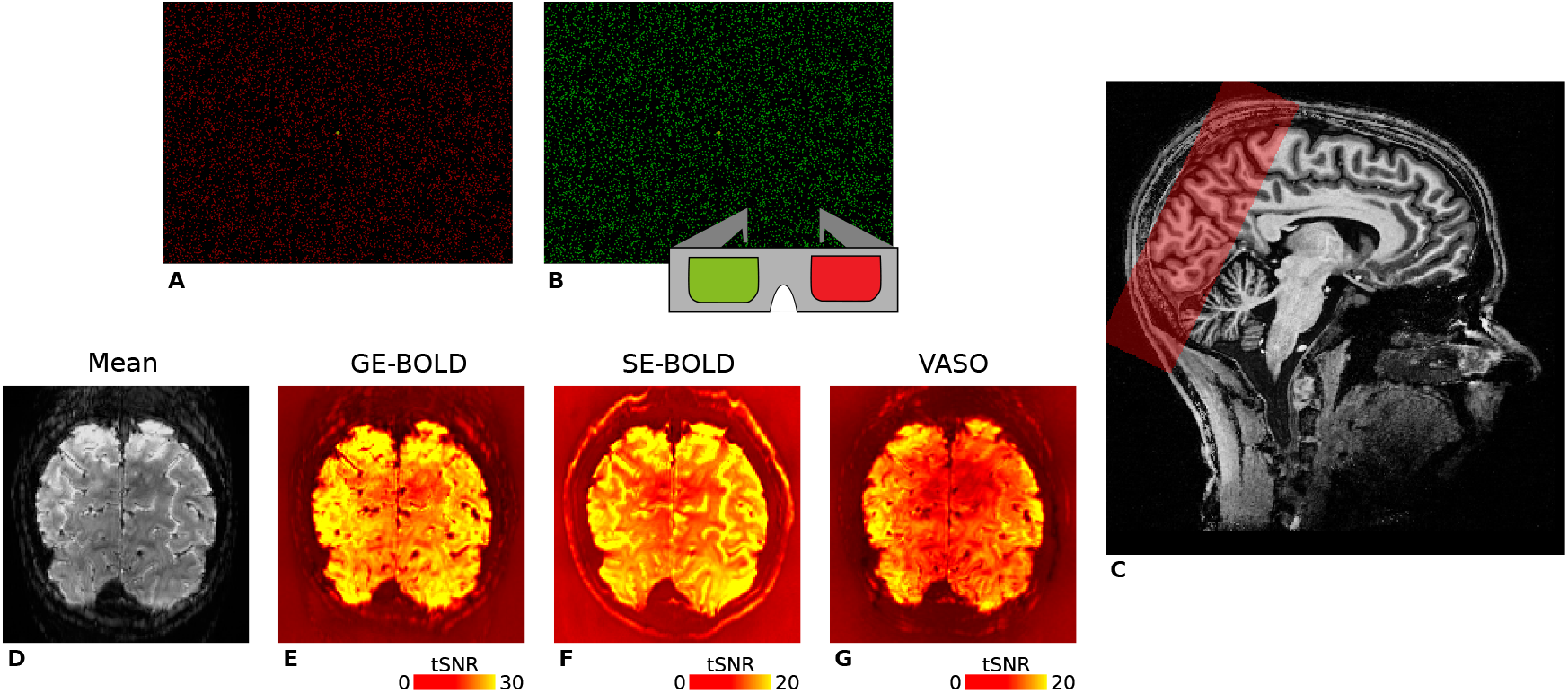
Illustration of the stimuli used for ocular dominance column mapping and representative fMRI data. For visual stimulation, we used red **(A)** and green **(B)** random dot stereograms (RDSs) that were viewed through anaglyph goggles by participants, respectively. Stimuli were based on Nasr, Polimeni, and Tootell, 2016 and enabled full field of view visual stimulation of the left or right eye in separate experimental blocks. RDSs formed the percept of an 8 × 6 checkerboard with independent sinusoidal movements in the horizontal direction of individual squares. **C** shows the spatial coverage of GE-BOLD acquisitions (red box) overlaid on a *T*_1_-weighted anatomical scan in sagittal view. Fewer slices were acquired for SE-BOLD and VASO sessions depending on specific absorption rate (SAR) limitations. For one representative participant (subject 1), the temporal mean of one GE-BOLD run and corresponding tSNR maps are shown in **D–G**. Note the different color scales.

### Visual stimulation

For the purpose of visual stimulation, an LCD projector (Sanyo PLC-XT20L) with custom-built focusing objective lens was used (refresh rate: 60 Hz, pixel resolution: 1024 × 768) that was positioned in the magnet room. To prevent interferences with the MR scanner, the projector was housed within a custom-built Faraday cage. The stimuli were projected onto a rear projection screen, mounted above the participants’ chest within the bore. Participants viewed the stimuli by means of a mirror attached to the head coil. In order to minimize scattered light reaching the participants’ eyes, the projection screen was surrounded by black felt, and all ambient lighting was turned off during data acquisition. This setup allowed visual stimulation within an approximate visual angle of 22° × 13° (width × height). Stimulus generation and presentation were carried out using the Psychophysics Toolbox (3.0.14, http://psychtoolbox.org/) (Brainard, 1997; Pelli, 1997; Kleiner et al., 2007) with GNU Octave (4.0.0, http://www.gnu.org/software/octave/).

#### ODC mapping

We used a block design with two experimental conditions that was previously reported in detail (Nasr, Polimeni, and Tootell, 2016; Haenelt et al., 2023), with the following minimal modifications for the current study. Every scanning session comprised ten runs, each lasting for 270 s. Within each run, a baseline period of 15 s was placed at the beginning and end, during which participants were presented with a uniform black background. The experimental protocol consisted of eight blocks, each lasting for 30 s, allowing four distinct stimulation periods targeting the left and right eye, respectively. The ordering of blocks was pseudorandomized. Throughout the runs, participants were instructed to maintain fixation on a central point (0.2° ×0.2°) and respond on a keypad when the fixation point changed its form (square or circle). Presented stimuli consisted of red or green random dot stereograms (RDS) (Julesz, 1971) shown on a black background (dot size: 0.1°, dot density: ~ 17%) that were viewed through custom-built anaglyph spectacles using Kodak Wratten filters No. 25 (red) and 44A (cyan), which enabled the stimulation of either the left or right eye in separate blocks, see ***Figure 1***. RDSs performed a horizontal sinusoidal movement (temporal frequency: 0.25 Hz, amplitude: 0.11°), and phases of dots were initialized to create the appearance of an 8 × 6 checkerboard with independent movement of squares. To reduce cross-talk between the eyes, the luminance of the dots was maintained at a low level (red through red filter: 3.1 cd/m^2^, red through cyan filter: 0.07 cd/m^2^, green through cyan filter: 5.7 cd/m^2^, green through red filter: 0.09 cd/m^2^). It is worth noting that the luminance of the green dots was approximately doubled rel-ative to red to ensure a similar excitation of cone photoreceptors for both colors (Dobkins, Thiele, and Albright, 2000).

#### Retinotopic mapping

To delineate the location of area V1, we employed a conventional phase-encoded paradigm (Sereno et al., 1995; Engel, Glover, and Wandell, 1997). Visual stimuli consisted of a flickering (4 Hz) black-and-white radial checkerboard restricted to a clockwise/anticlockwise rotating wedge (angle: 30°, temporal frequency: 1/64 Hz) or expanding/contracting ring (temporal frequency: 1/32 Hz) shown in separate runs. Each run presented 8.25 cycles of stimulation, with a baseline block of 12 s at the beginning and end of each run, in which a uniform gray background was shown. Runs lasted 552 s for the rotating wedge stimulus and 288 s for the moving ring stimulus. The mean luminance of the stimuli was set to 44 cd/m^2^. Throughout the run, participants were instructed to maintain fixation on a central point. No explicit task was given.

### Imaging

We used a whole-body MR scanner operating at 7 T (MAGNETOM 7 T, Siemens Healthineers, Erlangen, Germany) for measurements. The scanner was equipped with SC72 body gradients (maximum gradient strength: 70 mT/m; maximum slew rate: 200 mT/m/s). We used a single-channel transmit/32-channel receive head coil (Nova Medical, Wilmington, DE, USA) for RF signal transmission and reception. To optimize the transmit voltage over the occipital lobe, we always acquired a low-resolution transmit field map at the beginning of each scanning session using a sequence that exploits the ratio of consecutive spin and stimulated echoes (WIP-658).

For ODC mapping measurements, we acquired functional data with GE-BOLD, SE-BOLD, and VASO in different sessions. GE- and SE-BOLD data were acquired using a single-shot sequence with 2D echo planar imaging (EPI) readout (Feinberg et al., 2010; Moeller et al., 2010). For VASO measurements, we used a single-shot slice-selective slab-inversion (SS-SI) VASO sequence (Huber et al., 2014) with a 3D EPI readout (Poser et al., 2010). An oblique-coronal slab was imaged positioned over the occipital lobe. For all acquisition techniques, we used the following parameters: nominal voxel size = 0.8 mm isotropic, field of view (FOV) = 148 × 148 mm^2^, readout bandwidth (rBW) = 1182 Hz/px. For acceleration, we used GRAPPA = 3 with FLASH-based calibration (Talagala et al., 2016) and in-plane partial Fourier = 6/8 in the EPI phase-encoding direction, which resulted in an effective echo spacing of 0.33 ms. For GE- and SE-BOLD, we set the repetition time to TR = 3000 ms and used an echo time of TE = 24 ms and TE = 38 ms, respectively. The flip angle in GE-BOLD measurements was set to the Ernst angle FA = 77°, while in SE-BOLD, flip angles were set to 90° and 180° for excitation and refocusing pulses, respectively. For VASO measurements, we used an effective TR = 5000 ms, during which one image with (nulled) and one image without (not-nulled) blood nulling was acquired. Other parameters were the following: TE = 25 ms, TI = 1370 ms for the blood-nulling point, FA = 26°, 7.7% slice oversampling. 50 slices were acquired in GE-BOLD measurements that covered the whole stimulated area of V1. Due to specific absorption rate (SAR) constraints, the number of slices was limited for SE-BOLD and VASO measurements. For VASO, we acquired 26 slices. For SE-BOLD, we used the maximum number of allowed slices that varied between subjects and sessions and was between 16 and 29 slices.

A slightly modified GE-BOLD protocol was employed for retinotopy measurements, with the following parameters changed: voxel size = 1.0 mm isotropic, TR = 2000 ms, TE = 21 ms, FA = 68°, rBW = 1164 Hz/px, 40 slices.

For anatomical reference, we acquired a whole-brain anatomy using a 3D *T*_1_-weighted MP2RAGE sequence (Marques et al., 2010) with the following parameters: voxel size = 0.7 mm isotropic, TR = 5000 ms, TE = 2.45 ms, inversion times (TI1/TI2) = 900 ms/2750 ms with FA1/FA2 = 5°/3°, respectively, FOV = 224 × 224 × 168 mm^3^ (read × phase × partition), rBW = 250 Hz/px, GRAPPA = 2, partial Fourier = 6/8 (phase-encoding direction; outer loop). During online reconstruction on the scanner, a uniform *T*_1_-weighted image (UNI) was generated by combining data from both inversion times.

Protocols of all acquisitions are publicly available (https://osf.io/umnyr/).

### Data preprocessing

Functional time series from individual ODC mapping sessions were first subjected to motion correction to address within-run and between-run motion using SPM12 (v6906, https://www.fil.ion.ucl.ac.uk/spm/) with Matlab R2019b (MathWorks, Natick, MA, USA). Due to the used long stimulation periods and since transient time points were discarded in the analysis (see Pattern classification), no slice-timing correction was applied. In the case of VASO measurements, the time series were initially separated into individual time series for nulled and not-nulled images prior to motion correction. Motion correction was then independently applied to each of these time series. Final VASO time series were obtained by correcting the nulled time series for residual BOLD contamination. To achieve this, the motion-corrected nulled and not-nulled VASO time series were temporally upsampled onto a common grid using 3drefit from AFNI (19.1.05, https://afni.nimh.nih.gov/) (Cox, 1996), matching the effective temporal resolution of GE- and SE-BOLD measurements. Subsequently, the nulled time points were divided by the not-nulled time points to perform BOLD correction (Huber et al., 2014). All time series underwent then highpass filtering^1^ (cutoff frequency: 1/270 Hz), and a voxel-wise statistical analysis was performed for each session using a fixed-effects general linear model (GLM) as implemented in SPM12 with both experimental conditions as regressors convolved with the canonical hemodynamic response function (HRF). Note that GLM results were only used to visualize statistical maps and for the repeatability analysis (see Consistency of ocular dominance maps), while the main analysis was based on pre-processed fMRI time series.

The functional time series obtained from retinotopy measurements underwent similar pre-processing steps. However, prior to motion correction, each time series was corrected for different slice timings by voxel-wise temporal interpolation to a common time grid using 3drefit. Following motion correction, the time series were subjected to highpass filtering (cutoff frequency: 1/(3 × stimulus cycle period) Hz), which resulted in 1/192 Hz and 1/96 Hz for polar angle and eccentricity runs, respectively. The data from the first quarter stimulus cycle was discarded from further analysis. A voxel-wise Fourier transform was computed, and the signal at stimulus frequency was averaged from runs with opposite stimulus directions to compensate for the hemodynamic lag. The phase at stimulus frequency from polar angle runs was used to delineate the borders of V1.

To achieve registration between the reference anatomy and the functional time series without task, the anatomical image underwent an initial transformation to align with the functional space based on the scanner coordinate system. Only for registration, the mean functional image was bias field corrected (Tustison et al., 2010). Both images were then brain-masked and rigidly registered using ANTs (2.3.1, http://stnava.github.io/ANTs/). A similar procedure was employed for registering functional images from other sessions to the functional time series without task (between-session registration), except that a nonlinear registration was performed using the Symmetric Normalization (SyN) algorithm (Avants et al., 2008) implemented in ANTs.

The MP2RAGE (UNI) image was used for surface reconstruction of the cerebral cortex. Initially, the UNI image underwent bias field correction using SPM12. The corrected image was then fed into the recon-all pipeline in FreeSurfer (6.0.0, http://surfer.nmr.mgh.harvard.edu/) (Dale, Fischl, and Sereno, 1999; Fischl, Sereno, and Dale, 1999) with the hires flag to perform segmentation at the original voxel resolution (Zaretskaya et al., 2018). The brain mask was separately created based on the second inversion image of the MP2RAGE by using the SPM12 segmentation algorithm and excluding voxels in a binary mask that exceeded the tissue class threshold of 10% in all non-white matter (WM) and non-gray matter (GM) tissue classes. Subsequently, generated boundary surfaces of GM to WM and cerebrospinal fluid (CSF; pial boundary surface) were manually corrected, with particular attention given to the region surrounding the sagittal sinus. To counteract potential segmentation biases arising from basing FreeSurfer segmentation on the UNI image from the MP2RAGE, the resulting GM/WM boundary surfaces were shifted inward by 0.5 mm (Fujimoto et al., 2014). The final surfaces underwent smoothing using `mris_smooth` with 2 smoothing iterations implemented in FreeSurfer and were upsampled to an average edge length of approximately 0.3 mm.

Based on a computed registration between whole-brain anatomy and functional time series, boundary surfaces were transformed to the space of the reference EPI acquisition without task from the same session by applying the deformation field to surface vertices using linear interpolation. Functional images are spatially distorted in the phase-encoded direction due to the low bandwidth in this direction that leads to a sensitivity to variations in the main magnetic field. These distortions necessitate careful consideration (Jezzard and Balaban, 1995; Andersson, Skare, and Asburner, 2003), particularly when analyzing at the spatial scale of cortical layers.

We used the Gradient-Based Boundary (GBB) package (0.1.6, https://pypi.org/project/gbb/), which corrects the boundary surfaces by moving them to the GM/WM border found in functional images based on an iterative procedure, which is illustrated in ***Supplementary Figure 6***. To enhance the robustness of this method, we increased the GM/WM contrast in functional images following the method suggested in Fracasso, Petridou, and Dumoulin, 2016 that weights the magnitude image by its phase (both provided by the online reconstruction on the sanner) as conventionally practiced in susceptibility-weighted imaging methods^2^. For this purpose, the magnitude time series was corrected for motion using AFNI. Each image of the phase time series was individually phase unwrapped using the method by Abdul-Rahman et al., 2005 implemented in Nighres (1.2.0, https://pypi.org/project/nighres/) (Huntenburg, Steele, and Bazin, 2018), and computed motion parameters were subsequently applied to the unwrapped phase time series. The temporal mean of both magnitude and phase data was calculated, and the phase data underwent thresholding and normalization. Finally, the contrast of the magnitude data was enhanced by assigning weights to each voxel based on the contrast-reversed phase data.

Nine equidistant surfaces were computed and positioned between boundary surfaces^3^. This resulted in 11 cortical layers for subsequent analyses.

For sampling data onto reconstructed surfaces, surfaces were first moved into the space of individual functional sessions based on the computed registration. Subsequently, the functional data were sampled onto the surface mesh using linear interpolation.

### Pattern classification

We used a linear support vector machine (SVM) algorithm for pattern classification from single time points of motion-corrected and detrended functional time series. Each ODC mapping session and each cortical depth was analyzed independently. For classification, functional time series were first sampled onto a cortical layer. One run contained 90 time points, and 10 runs were acquired per session. All time points from the baseline condition were discarded. Additionally, the first two time points from each experimental condition were discarded from further analysis to omit contamination from transient effects of the hemodynamic response during classification. This resulted in 64 time points per run, evenly divided between left and right eye stimulation. Sampled time series were then standardized and divided into a training data set (9 runs, 576 time points) and a test data set (1 run, 64 time points).

Feature selection was performed by only considering time series data from locations within V1 that were present in the FOV of all functional sessions. Based on the training data, we further used an *F*-test implemented in the scikit-learn library (1.2.0, https://scikit-learn.org/) (Pedregosa et al., 2011), specifically `sklearn.feature_selection.f_classif`, to select the vertices whose time series strongest correlated with the experimental paradigm. We used the training data averaged across cortical depth to select the same features across cortical depth. The top 200 vertices with the highest correlation were chosen for further analysis. The number of selected vertices was determined by selecting less features than samples to decrease the chances of overfitting as similarly done in Haynes and Rees, 2005b.

For classification, we used the SVM implementation `sklearn.svm.SVC` with fixed regularization term *C* = 1 that is based on the `libsvm` library (Chang and Lin, 2011). This method was repeated for all possible splittings of training and test data sets using a leave-one-run-out cross-validation procedure to estimate mean prediction accuracies.

## Results

### Topography of ocular dominance columns

***Figure 2*** shows ocular dominance column maps (contrast: left eye > right eye) for a representative participant sampled at mid-cortical depth. Maps from single participants can be found in ***Supplementary Figure 1***–***Supplementary Figure 5. Figure 2A*** shows the average activation map across two GE-BOLD sessions. Some features can be seen that are expected from ODCs: (1) V1 shows a fine-scale pattern. (2) The pattern is constrained to area V1. (3) Around the approximate location of the horizontal meridian, columns are oriented more in parallel to both vertical meridians (V1/V2 border) (LeVay, Hubel, and Wiesel, 1975). This is the expected topography as depicted in (Adams, Sincich, and Horton, 2007; Adams and Horton, 2009).

**Figure 2.**
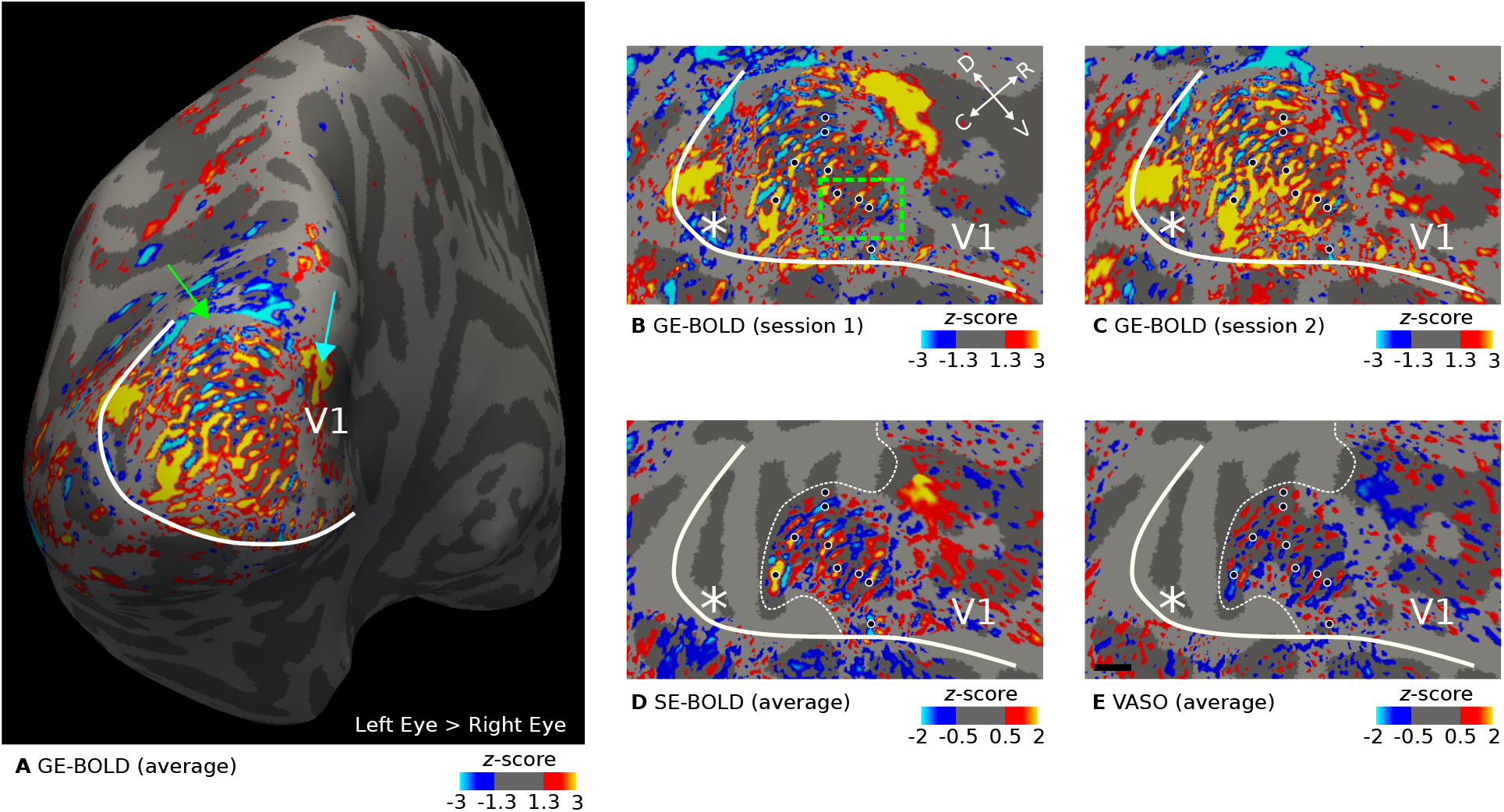
Representative maps of ocular dominance columns (ODCs). Thresholded activation maps (contrast: left eye > right eye) are shown for the left hemisphere of one representative participant (subject 1). Data were sampled at mid-cortical depth. In **A**, the contrast from GE-BOLD sessions (average across two sessions) is shown on the inflated surface. Several columns confined to V1 can be identified. The green arrow points to a small location (gray area) outside of the imaging field of view. **B–C** show the contrast from single GE-BOLD sessions on the flattened surface. The similar appearance of both maps illustrates the consistency of the columnar pattern across sessions conducted on different days. **D–E** show the contrast from SE-BOLD and VASO sessions (average across sessions). Due to the reduced number of slices, the area around the foveal representation was not covered (see the white dotted line that outlines the covered area). A similar ocular dominance pattern can be seen in all maps (see black dots with white outline for reference). Note that VASO has an inverted contrast compared to BOLD. The white line shows the representation of the vertical meridian (V1/V2 border) that was based on a separate retinotopy measurement. White asterisks indicate the location of the foveal representation. The black line in **E** shows a scale bar (5 mm). Maps from all participants can be found in ***Supplementary Figure 1***–***Supplementary Figure 5***.

The blind spot is a further distinctive monocular region of V1 (Tootell et al., 1998). Due to the lack of photoreceptor cells on the optic disc of the retina where the optic nerve bundles and passes through, there is an oval area in V1 on the contralateral hemisphere that is solely “filled” by the response from the ipsilateral eye. In ***Figure 2A***, there is a spatially extended response from the ipsilateral eye at the anterior end of the stimulated area (see cyan arrow in ***Figure 2A***), which could be the blind spot representation on this hemisphere. However, due to the limited visual field in our experiment, we did not expect to have covered the blind spot region, which should be found at around 15° eccentricity (Tootell et al., 1998). Therefore, we assume that this response is of vascular origin or a response that was elicited by the border of the stimulus. This region was carefully left out in the decoding analysis.

We cannot exclude the possibility that some columns merged due to idiosyncrasies in local vasculature, which might explain the appearance of some broader activation clusters in V1. Further analyses of possible mechanisms would be compelling but is outside of the scope of the current study. But interestingly, these clusters were repeatable across sessions, as can be seen when comparing ***Figure 2B*** and ***Figure 2C*** that show GE-BOLD activation maps from single sessions. The comparison also indicated the overall high consistency of ODCs between sessions. This was also confirmed by the fact that the pattern remained stable after averaging, as shown in ***Figure 2A***. A more quantitative repeatability analysis is given in the next section (Consistency of ocular dominance maps). Black dots are displayed to aid comparison of ODC patterns between maps.

***Figure 2C*** and ***Figure 2D*** show the average activation maps across sessions for SE-BOLD and VASO, respectively. Due to SAR constraints (see Materials and methods, fewer slices were acquired for SE-BOLD and VASO. Coverage boundaries are outlined by white dotted lines. However, within the imaged region, a similar ODC pattern can be identified at the expense of overall reduced signal strength.

For the inset presented in ***Figure 2B, Figure 3*** illustrates the unthresholded contrast sampled at different cortical depths. It can be seen that certain columns display consistent activation through the cortical ribbon, suggesting a degree of columnar stability.

**Figure 3.**
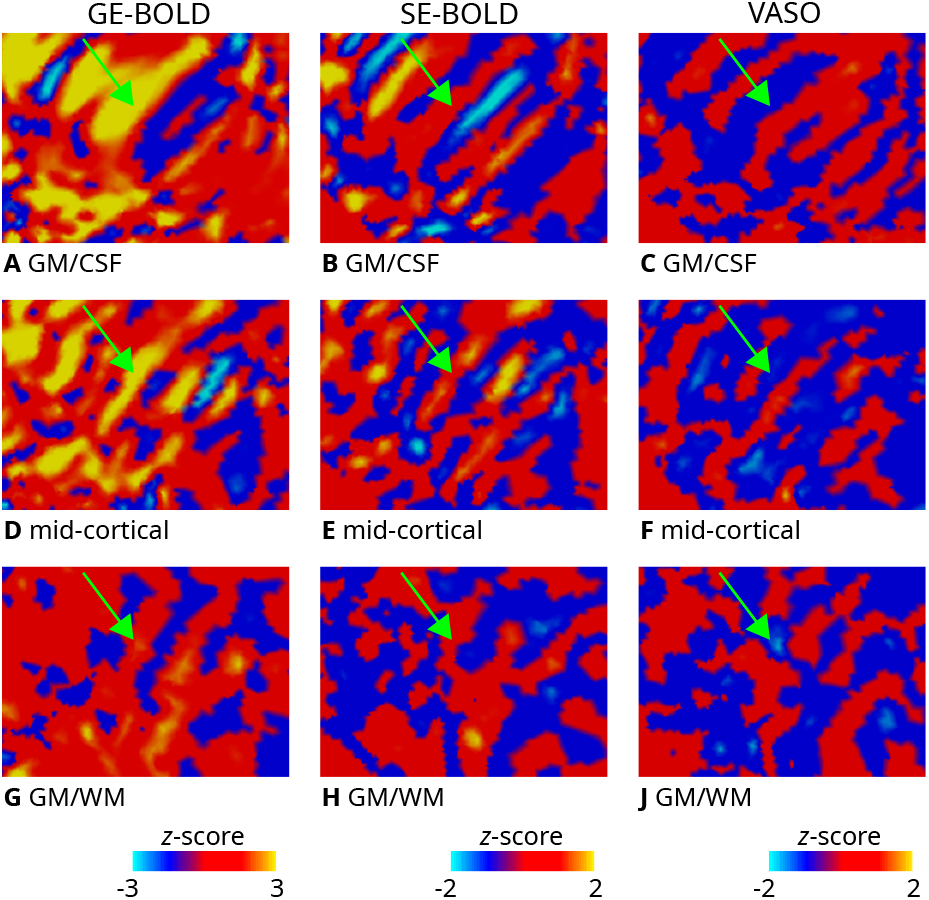
Zoomed view of ODC maps. Unthresholded activation maps (contrast: left eye > right eye; average across two sessions) are shown for the left hemisphere of one representative participant (subject 1). The section shown corresponds to the inset (green rectangle) defined in ***Figure 2B***. Data were sampled on the flattened GM/CSF **(A–C)**, mid-cortical **(D–F)**, GM/WM **G–J** boundary surfaces for GE-BOLD (left column), SE-BOLD (middle column), and VASO (right column), respectively. Despite lower SNR of SE-BOLD and VASO, some similar patterns can be identified across contrasts and cortical depth (see green arrow). Note that VASO has an inverted contrast compared to BOLD and different color scales were used.

### Consistency of ocular dominance maps

We quantified the repeatability of ocular dominance maps between sessions. For this purpose, we computed Spearman’s rank correlation coefficient between *z*-scores (contrast: left eye > right eye) restricted to mid-cortical depth from both sessions of each acquisition method. In the analysis, only vertices within V1 were considered that were located within the FOV of all sessions.

***Figure 4*** shows scatter plots for one representative participant. Spearman’s rank correlation coefficient and the corresponding *p*-value are stated in the figures, which demonstrates the repeatability of elicited responses across sessions. The *p*-value was determined by permutation testing. A null distribution was created by computing the correlation coefficients between data from the first session and spatially shuffled data from the second session (*n* = 10, 000). The *p*-value was then calculated as the fraction of the null distribution greater or smaller than the computed statistics with unshuffled maps. Considering the non-independency of data from neighboring vertices, we used only a fraction of randomly chosen 10% of vertices for the analysis (Nasr, Polimeni, and Tootell, 2016).

**Figure 4.**
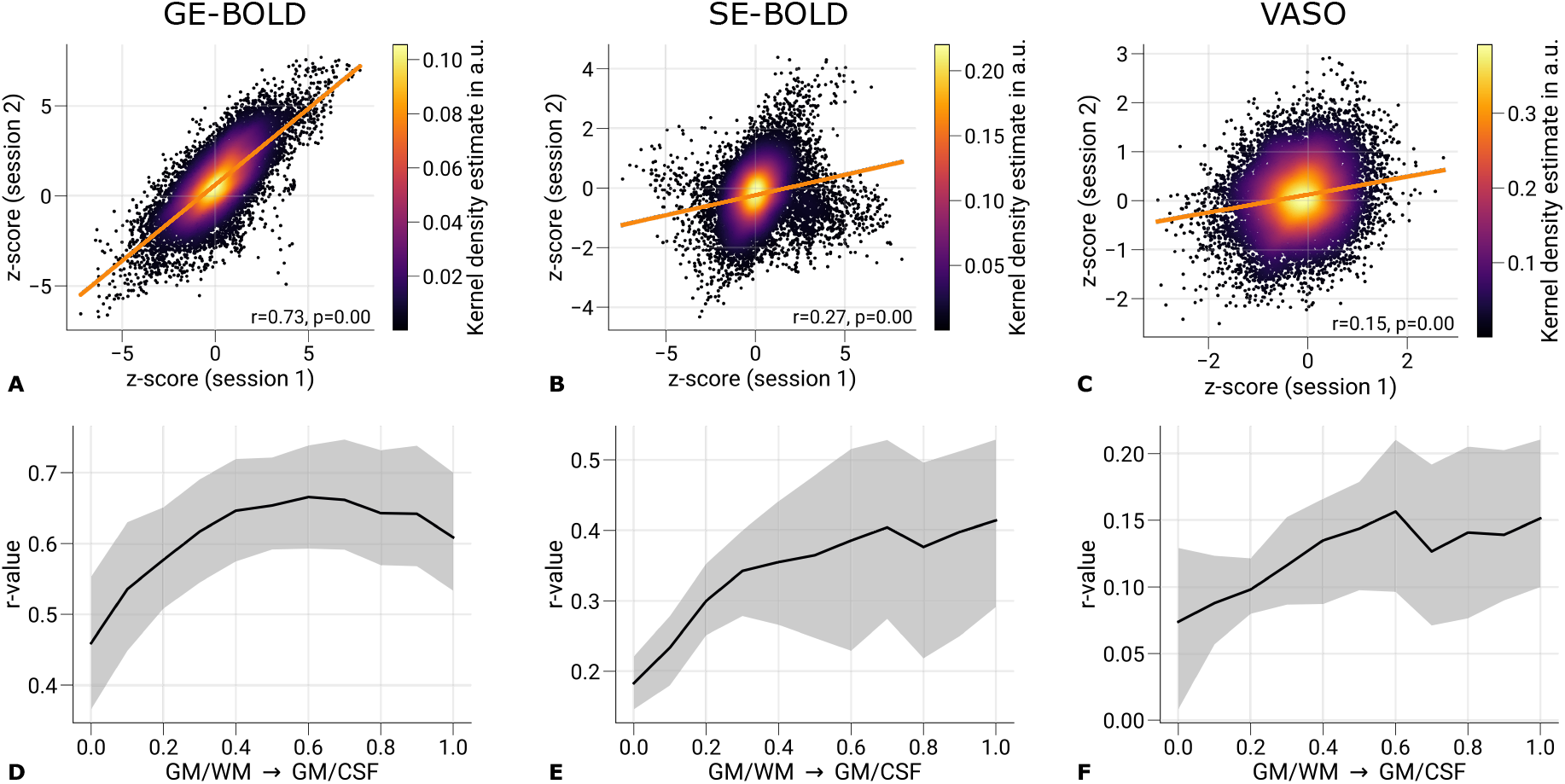
Repeatability of ODC maps across scanning sessions. Scatter plots with kernel density estimation illustrate the consistency of activation maps (contrast: left eye > right eye) across GE-BOLD **(A)**, SE-BOLD **(B)**, and VASO **(C)** scanning sessions for one representative participant (subject 1). Only data from V1 sampled at mid-cortical depth were used. The regression line is shown as an orange line. Spearman’s rank correlation coefficients and corresponding *p*-values are stated next to the plots. Statistical significance was determined by permutation testing (n = 10,000). Due to the spatial covariance of data from neighboring vertices, only randomly selected 10% of all data points were used for significance testing. In **D–F**, the mean correlation is shown across cortical depth. Black lines indicate the mean across participants and scanning sessions. The gray area demarcates the bootstrap 95% confidence interval (n = 1,000). See ***Table 1*** for the results of the correlation analysis from all participants.

We acknowledged the variability *σ* of the estimated *p*-value due to the finite size of generated null distributions. A correction was applied by modeling the variability by the variance of a binomial distribution σ^2^ = *np*(1−*p*) and adding an upper bound of 3σ to the number of samples exceeding the test statistics (Burt et al., 2020). A corrected *p*-value of < 0.05 was considered statistically significant.

**Table 1.** Repeatability of ODC maps across scanning sessions for single participants. Spearman’s rank correlation coefficients and corresponding *p*-values are shown to illustrate the consistency of activation maps (contrast: left eye > right eye) between scanning sessions for single participants. Only data from V1 sampled at mid-cortical depth were used. Statistical significance was determined by permutation testing (n = 10,000). Due to the spatial covariance of data from neighboring vertices, only randomly selected 10% of all data points were used for significance testing.

|  | GE-BOLD |  | SE-BOLD |  | VASO |  |
| --- | --- | --- | --- | --- | --- | --- |
|  | Correlation coefficient (*r*) | *p*-value | Correlation coefficient (*r*) | *p*-value | Correlation coefficient (*r*) | *p*-value |
| Subject 1 | 0.623 | < 0.001 | 0.219 | < 0.001 | 0.129 | < 0.001 |
| Subject 2 | 0.634 | < 0.001 | 0.185 | < 0.001 | 0.049 | < 0.05 |
| Subject 3 | 0.755 | < 0.001 | 0.493 | < 0.001 | 0.186 | < 0.001 |
| Subject 4 | 0.586 | < 0.001 | 0.418 | < 0.001 | 0.167 | < 0.001 |
| Subject 5 | 0.643 | < 0.001 | 0.379 | < 0.001 | 0.132 | < 0.001 |

***Figures 4D–F*** illustrate the correlation between sessions across cortical depth. All plots show an increase in correlation toward the pial surface, which matches the typically seen increase in signal changes in BOLD acquisitions. However, correlation coefficients decrease again in upper layers in ***Figure 4D***. This might be explained by overall higher temporal variability in upper cortical layers caused by multiple sources, e.g., brain pulsatility, which lead to dynamic partial volume changes with the high-intensity CSF signal (Polimeni et al., 2010b).

Overall, the correlation coefficients were relatively low. However, this outcome is expected given that the analysis included all V1 vertices rather than a subset with most strongly activated clusters as in the main analysis, which could have artificially inflated the correlation estimates. The lower correlations thus partly reflect the fact that ODCs were not uniformly resolvable across V1, with stable columnar patterns observed only in a subset of locations, as illustrated in ***Figure 2***. Whether these more consistent regions are driven by vascular or neuronal factors remains an open question and is beyond the scope of the present study.

***Table 1*** summarizes the correlation results across all participants.

### Univariate contrasts across cortical depth

***Figure 5*** shows the strength of cortical responses by plotting the percent signal changes of left and right eye stimulation across cortical depth. The mean across participants and sessions and the corresponding 95% bootstrap confidence interval are shown. Red lines (solid and dashed) depict the mean response for single sessions, demonstrating the repeatability of cortical profiles.

**Figure 5.**
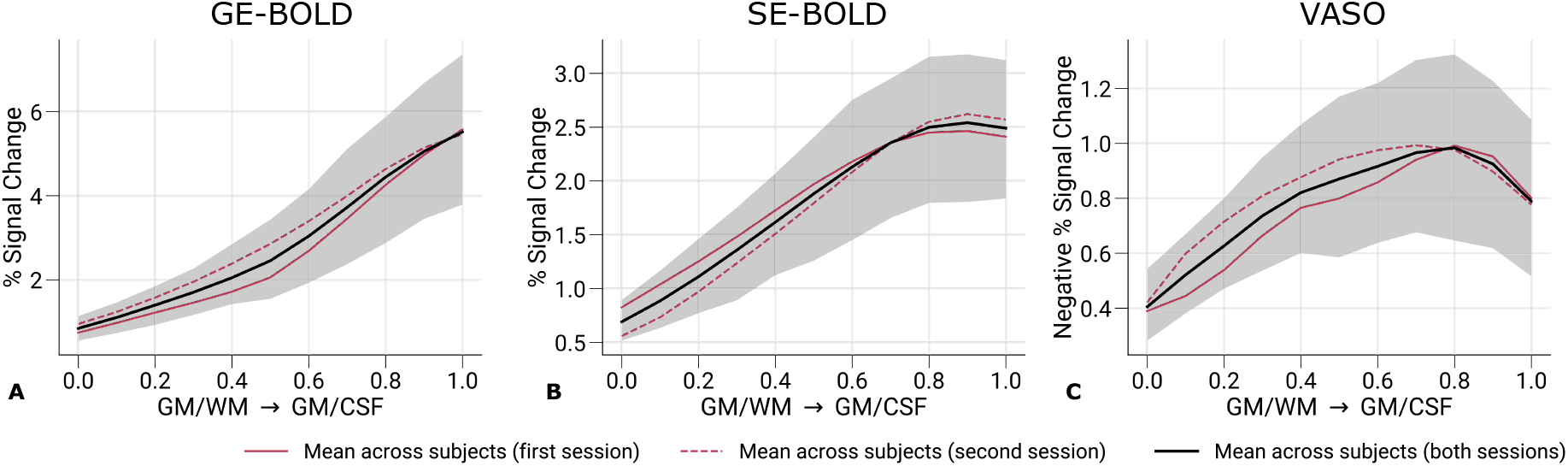
Percent signal changes across cortical depth. Mean percent signal changes (contrast: left eye and right eye > baseline) for GE-BOLD **(A)**, SE-BOLD **(B)**, and VASO **(C)** are shown across cortical depth. Red solid and dashed lines show the mean across participants from the first and second session, respectively. Black lines indicate the mean across participants and scanning sessions. The gray area demarcates the bootstrap 95% confidence interval (n = 1,000). Only data points (n = 200) were used that were also selected for the decoding analysis. Note that we inverted the *y*-axis in **C** for consistency with **A** and **B**. Mean percent signal changes across cortical depth with all V1 data can be found in ***Supplementary Figure 7***. Percent signal change curves from single participants can be found in ***Supplementary Figure 8***.

We used the same vertices that were included in the classification analysis after feature selection. As expected, GE-BOLD signal changes were overall larger than SE-BOLD and VASO. Note that signal changes for VASO, which has a negative relationship with CBV changes, were inverted for visual purposes.

Across cortical depth, both GE- and SE-BOLD showed a steady increase toward the pial surface, most likely reflecting draining vein contributions to the signal (Polimeni et al., 2010a; Markuerkiaga, Barth, and Norris, 2016). The VASO signal profile was more restricted to GM and shows a peak within GM. But an overall trend toward the pial surface could be seen as well. In ***Supplementary Figure 7***, cortical profiles of signal changes across participants are shown with all V1 vertices included. In these plots, VASO shows a more pronounced peak within GM. However, due to the averaging across more data points, V1 vertices that were not activated and therefore only contain noise contributions were included, which led to a general decrease of percent signal changes from all acquisition techniques. This suggests the hypothesis that the often seen reduced signal changes at the pial surface and pronounced peak within gray matter for SS-SI VASO may partly be driven by inclusion of pure signal noise. ***Supplementary Figure 8*** further illustrates cortical profiles of signal changes from single participants, demonstrating the variability between participants in our study.

### Decoding accuracies across cortical depth

***Figure 6*** shows mean prediction accuracies across cortical depth from the pattern classification analysis. An independent classification was performed for each cortical depth with features selected from the mean response across cortical depth. Black lines indicate the mean across participants and sessions with the corresponding 95% bootstrap confidence interval. Red lines depict mean prediction accuracies from single sessions. ***Supplementary Figure 9*** further illustrates prediction accuracies from single participants.

**Figure 6.**
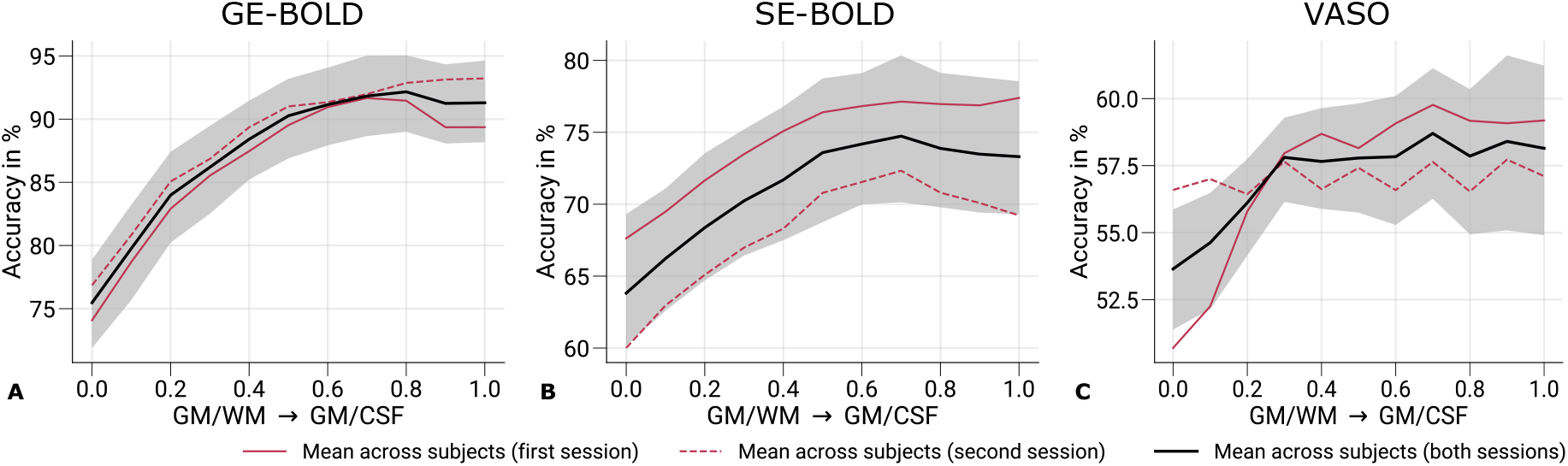
Prediction accuracies across cortical depth. Mean prediction accuracies (prediction of the stimulated eye) for GE-BOLD **(A)**, SE-BOLD **(B)**, and VASO **(C)** are shown across cortical depth. Red solid and dashed lines show the mean across participants from the first and second session, respectively. Black lines indicate the mean across participants and scanning sessions. The gray area demarcates the bootstrap 95% confidence interval (n = 1,000). In **A–C**, data were significantly different (*p* < 0.05) from a 50% chance level at each cortical depth. The *p*-value was determined by bootstrapping (n = 1,000) and corrected for multiple comparisons of individual layers (FDR correction using the Benjamini and Hochberg procedure). Prediction accuracy curves from single participants can be found in ***Supplementary Figure 9***.

With all acquisition techniques, the eye-of-origin could be decoded with statistical significance at all cortical depths (chance level: 50%, *p*-value determined by bootstrapping). Among acquisition techniques, GE-BOLD showed the highest prediction accuracies. Furthermore, prediction accuracies increased toward the pial surface, mirroring the increase of univariate responses across cortical depth as shown in the previous section (Univariate contrasts across cortical depth). However, prediction accuracies did not show a steady increase compared to signal change profiles but saturated around mid-cortical depth, more resembling the cortical profile from the repeatability analysis (Consistency of ocular dominance maps) A similar behavior could be seen for SE-BOLD with an overall reduced level of prediction accuracies.

Since VASO encodes volumes without blood nulling that are purely BOLD-weighted in addition to time points with blood nulling, we also used the not-nulled time points for decoding the eye-of-origin, which is shown in ***Supplementary Figure 10***. Overall, a similar profile to ***Figure 6A*** can be seen with general lower decoding accuracies that is most probably related to the lower temporal efficiency of the VASO measurements due to the longer volume TR.

From a neuronal perspective, one would have expected highest eye-of-origin decoding in deeper cortical layers since thalamocortical projections from the LGN primarily enter in layer 4C of V1 (Nieuwenhuys, Voogd, and Huijzen, 2008), which is located slightly below mid-cortical depth, see (Weber et al., 2008; Oga, Okamoto, and Fujita, 2016). Despite the anticipated higher laminar specificity of VASO, the decoding profile also showed a large resemblence to the profiles obtained with GE- and SE-BOLD. This suggests that remaining macrovascular contributions also limit the laminar specificity in VASO.

To better understand the potential impact of the feature selection process, we also conducted exploratory analyses by changing the cortical depth at which the feature selection process, which is presented in ***Supplementary Figure 11*** and ***Supplementary Figure 12***. In the main analysis, features were selected based on the training data averaged across all cortical depths, with the rationale of preserving the columnar organization by applying the same features set across cortical depth. Interestingly, ***Supplementary Figure 12*** reveals that when feature selection is confined to deeper cortical layers, a peak in decoding performance appears to emerge slightly below mid-cortical depth as expected for monocular thalamocortical input. This change in decoding pattern across cortical depth is more prominent in VASO compared to GE- and SE-BOLD. These findings suggest that excluding superficial layers—more susceptible to physiological noise and large draining veins—during feature selection may help uncover the enhanced laminar specificity inherent to VASO. Nonetheless, these results should be interpreted with caution, and further systematic investigations are required to confirm this effect, which lies beyond the scope of the present study.

## Discussion

In this study, we used high-resolution fMRI at sub-millimeter resolution to map ODCs in human V1 and decoded the eye-of-origin from pre-processed fMRI time courses. High-resolution imaging has previously characterised the depth profile of ODCs with GE-BOLD (Hollander et al., 2021) and VASO (Akbari et al., 2023). Building on this work, we directly compared the laminar specificity of eye-of-origin decoding across three contrasts—GE-BOLD, SE-BOLD, and VASO.

Early MVPA studies showed that eye-of-origin and orientation information could be decoded from V1 even with conventional resolution (3 × 3 × 3 mm^3^) (Kamitani and Tong, 2005; Haynes and Rees, 2005a; Haynes and Rees, 2005b). Those findings sparked debate about whether the classifiers exploited columnar signals or coarse-scale biases (Boynton, 2005; Beeck, 2010; Swisher et al., 2010; Gardner, 2010; Shmuel et al., 2010; Kriegeskorte, Cusack, and Bandettini, 2010; Chaimow et al., 2011; Misaki, Luh, and Bandettini, 2013). Because LGN inputs terminate monocularly in layer 4C and become increasingly binocular after intracortical processing (Wandell, 1995), the cortical depth profile of decoding accuracy can help to disambiguate these sources.

Our sub-millimeter fMRI acquisitions allowed us to sample the functional signal across cortical depth with sufficient resolution to study laminar differences. By tracking decoding performance as a function of depth, we assessed how much eye-of-origin information is available at each lamina and under each contrast. Finally, since macrovascular draining might act as a spatial-temporal filter that redistributes columnar signals to coarser scales (Kriegeskorte, Cusack, and Bandettini, 2010), depth-dependent decoding also potentially provides a means to distinguish microvascular from macrovascular contributions to the patterns exploited by the classifier.

As a prerequisite, we demonstrated robust in vivo mapping of ODCs across all acquisition methods, as shown in ***Figure 2*** (see ***Supplementary Figure 1***–***Supplementary Figure 5*** for activation maps of all participants). The observed activation patterns were consistent across imaging sessions and aligned well with ODC topographies previously reported in postmortem histological studies (Adams, Sincich, and Horton, 2007; Adams and Horton, 2009). In addition to the expected fine-scale columnar structures, some activation maps exhibited larger, coarser clusters that may reflect vascular contributions, particularly from regions dominated by larger draining veins. However, pin-pointing the exact source of these larger clusters is beyond the scope of the present study. ***Figure 3*** further illustrates the columnar nature of these patterns across cortical depth. Note that the consistency of the cortical-depth dependent ODC reponse was also shown in earlier results (Haenelt et al., 2019).

Overall, both SE-BOLD and VASO produced lower signal changes and exhibited increased noise levels, consistent with their inherently lower SNR. Despite these limitations, a subset of ODCs could be reliably mapped across sessions for all acquisition types. This reduced SNR was reflected in the repeatability analysis shown in ***Figures 4A–C*** and ***Table 1***. The session-to-session correlations of ODC maps were highest for GE-BOLD, followed by SE-BOLD and VASO. Depth-resolved visualizations of inter-session correlation (***Figures 4D–F***) revealed increasing repeatability toward the pial surface, likely driven by stronger signal contributions from macrovasculature in upper layers. Notably, for GE-BOLD (***Figure 4A***), the correlation did not increase monotonically across cortical depth but instead dropped in the outermost layers, likely due to higher signal variability near the CSF boundary (Polimeni et al., 2010b).

The MVPA analysis revealed that eye-of-origin information could be reliably decoded from fMRI time series across cortical depth for all acquisition methods, see ***Figure 6***. Decoding performance was highest for GE-BOLD, followed by SE-BOLD and VASO. These decoding profiles closely mirrored the patterns observed in the repeatability analysis, underscoring the critical role of signal-to-noise ratio (SNR) in classifier performance. Notably, decoding accuracy peaked around mid-cortical depth, in contrast to the monotonic increase in signal amplitude across depth observed in univariate analyses shown in ***Figure 5***. As discussed earlier, if the classifier primarily relied on laminar-specific information, we would expect a peak in deeper layers, particularly around layer 4C, where monocular input is most segregated. The absence of such a peak suggests that laminar specificity is limited across all acquisition types (but see further below for a discussion on the role of feature selection).

For VASO measurements, we initially expected to see increased laminar specificity by enhanced responses in deeper layers. A recent ODC mapping study by Akbari et al., 2023 indeed reported a peak in deeper layers in univariate response profiles from data sampled in V1. Differences between studies, including experimental design, acquisition parameters, or analysis choices, may underlie these discrepancies but cannot be completely resolved in this study. One possible factor, however, might be differences in the definition of regions of interest (ROIs). In our study, ROIs for univariate cortical profiles in ***Figure 5*** were based on the same feature selection process as for the main decoding analysis, which might have biased voxel selection toward regions with increased macrovascular contributions and elevated SNR. For example, ***Supplementary Figure 7*** shows univariate profiles with all V1 voxels included, where, the VASO response peaks closer to the mid-cortical depth. However, including all voxels introduces additional noise, particularly in superficial layers where partial volume effects with CSF are more pronounced (Polimeni et al., 2010b; Pfaffenrot et al., 2021).

Higher spatial resolution is expected to decrease this effect. Interestingly, a recent study by Feinberg et al., 2022 employed GE-BOLD and VASO acquisitions with an isotropic voxel size of 0.4 mm, i.e., an 8-times smaller voxel volumes compared to the current study, which showed a pronounced peak in deeper cortical layers in V1 for binocular visual stimulation. In addition, a second peak was observed in the upper layers. When considering feedforward thalamocortical input to V1, the deeper peak likely reflects input to layer 4, while the superficial peak may result from cortico-cortical processing or residual contributions from draining veins. Thus, the double-peak profile observed by Feinberg et al., 2022 may reflect a combination of neuronal and vascular origins.

In the main decoding analysis, feature selection was based on the mean cortical response. This ensured that the same vertices were selected across cortical depth, acknowledging the columnar topography of ODCs in V1. However, this approach may bias selection toward regions with higher SNR, which are also more likely to contain macrovascular contributions. Feature selection based on data further away should decrease these contributions. To address this, we conducted an additional analysis where we selected features solely from data sampled at the GM/WM, mid-cortical, GM/CSF surface, respectively, and independently for each cortical depth. The resulting univariate and decoding profiles are shown in ***Supplementary Figure 11*** and ***Supplementary Figure 12***, respectively. These results highlight the influence of feature selection on the observed profiles. For instance, univariate reponses in ***Supplementary Figure 11*** show that GE-BOLD shows a steady increase toward the pial surface irrespective of the feature selection process. However, SE-BOLD and VASO only exhibit a steady increase if feature selection is based on the GM/CSF surface. This behavior is also mimicked in decoding profiles shown in ***Supplementary Figure 12***. Interestingly, VASO shows a peak below mid-cortical depth, which does not coincide with the GM/WM surface, when feature selection is based on the GM/WM surface, further away from macrovascular contributions at the pial surface. Conversely, when feature selection is based on the GM/CSF surface, VASO shows a peak above mid-cortical depth. In case of independent feature selection for each cortical depth, this sums up to the resemblence of a double-peak (see ***Supplementary Figure 12M***) similar to Feinberg et al., 2022. The deeper peak corresponds to the approximate location of layer 4C (Palomero-Gallagher and Zilles, 2019) (relative cortical depth of 73%). This might hint to increased laminar specificity inherent in the VASO signal that might be exploited by the classifier, but also shows the dependence on the chosen feature selection process. However, due to the low sample size, this eploratory analysis prohibits detailed analysis and awaits further study. Future studies might want to reproduce and locate the exact cortical depth of the peak by combining using myelin-sensitive MRI acquisitions (Stüber et al., 2014; Trampel et al., 2019; Weiskopf et al., 2021) to locate the stria of Gennari (Trampel, Ott, and Turner, 2011; Fracasso et al., 2016) as a reference depth, see e.g. (Koopmans, Barth, and Norris, 2010; Huber et al., 2021).

Another methodological factor in our study is the arbitrary choice of the number of features used for classification. The main decoding analysis was restricted to 200 features (vertices). To investigate the effect of feature number on decoding performance, we conducted an additional analysis in which prediction accuracies were computed as a function of the number of selected vertices [1, 2, …, 500]. Results are shown in ***Figure 7***. It can be seen that only a few voxels were necessary to decode the eye-of-origin, which was similarly found for orientation decoding (Haynes and Rees, 2005a). GE- and SE-BOLD show a consistent trend across number of features with saturation at mid-cortical depth for prediction accuracies (***Figures 7A–B***) and steady increase of univariate responses toward the pial surface (***Figures 7D–E***). In contrast, VASO exhibited more variable patterns (***Figure 7C***) and showed a tendency for increased decoding accuracies at deeper layers. Corresponding univariate responses (***Figure 7F***) also peaked at mid-depth, which got more pronounced with increased number of features (cf. with univariate profile based on all V1 voxels shown in ***Supplementary Figure 7***). Additionally, ***Supplementary Figure 13*** illustrates decoding results using depth-specific feature selection at varying feature numbers. While GE- and SE-BOLD results remained stable, an apparant peak emerged at deeper layers for VASO. However, due to the limited dataset, these trends require further statistical validation.

**Figure 7.**
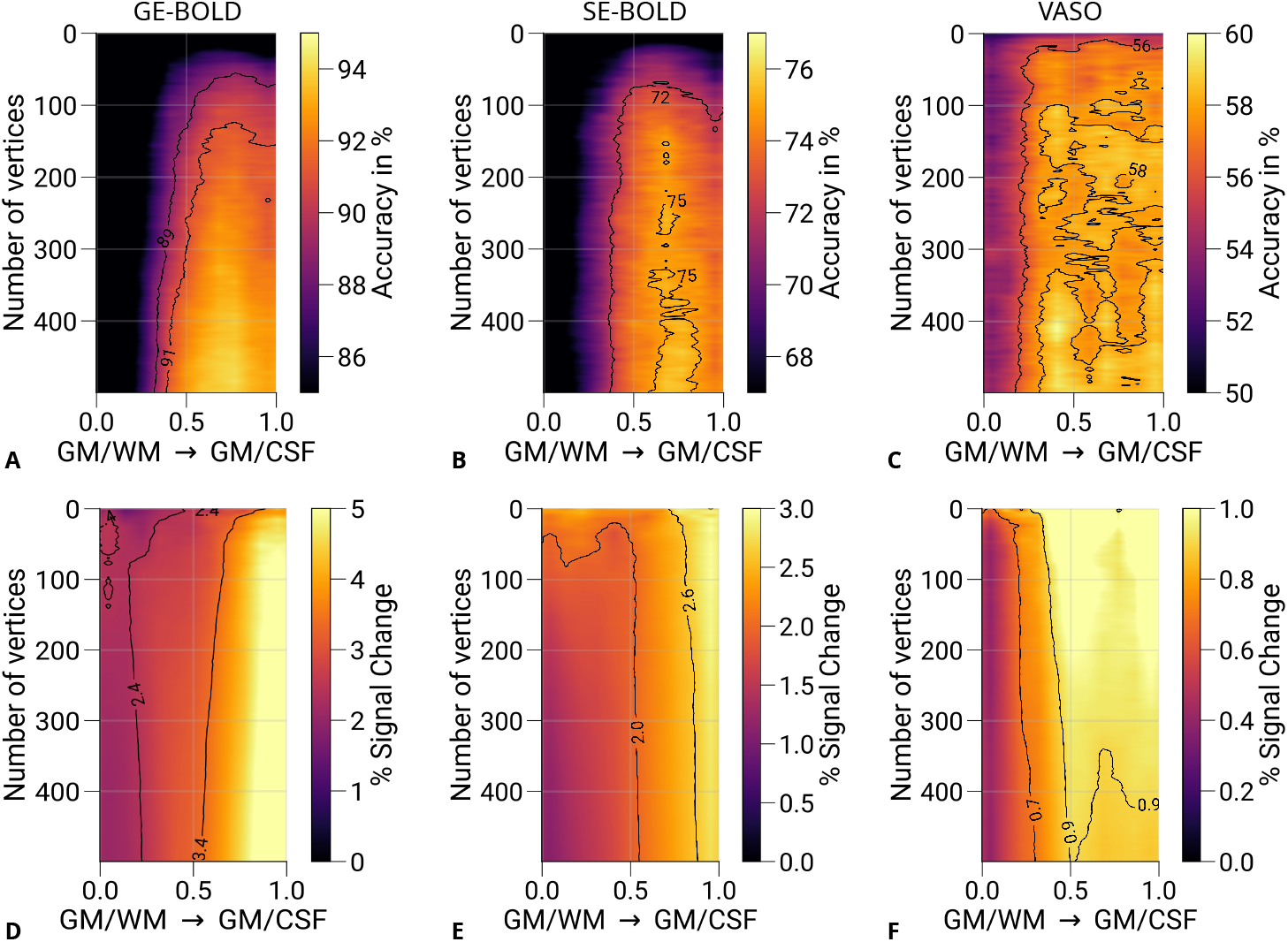
Prediction accuracies and percent signal changes for different number of features. Mean prediction accuracies (prediction of the stimulated eye) for GE-BOLD **(A)**, SE-BOLD **(B)**, and VASO **(C)** are shown for a varying number of features (vertices) across cortical depth. Note that 200 vertices were used for the principal analysis (see ***Figure 6***). **D–F** show corresponding percent signal changes (left eye and right eye > baseline) using the same data points also selected in the decoding analysis. Both prediction accuracies and percent signal changes appear to peak closer to the GM/WM boundary compared to GE- and SE-BOLD, respectively. Isolines are shown as black lines. For visualization purposes, images were slightly smoothed with a Gaussian kernel.

The interpretation of the laminar profile is built on the assumption that the monocular feed-forward information is exploited in V1, which is encoded at the fine-grained level of ODCs. Note that the larger monocular regions in V1, like the blind spot (Tootell et al., 1998) and the temporal monocular crescent (Nasr et al., 2020), were not covered in our experiment due to the limited field of view. However, we cannot exclude that other features besides ocularity might have contributed to the successful eye-of-origin decoding. Therefore, we conducted an additional analysis, in which we decoded the stimulated eye from cortical areas outside of V1 that are known not to be driven by monocular input. ***Figure 8*** shows cortical profiles of prediction accuracies from GE-BOLD data (200 vertices) sampled in the secondary visual cortex (V2) and the tertiary visual cortex (V3), respectively. V2 and V3 were further divided into two halves (*a*: half closer to V1, *b*: half further away from V1). The stimulated eye could be decoded in both V2 and V3 across cortical depth, but with overall decreased decoding performance compared to ***Figure 6A***. Furthermore, a similar increase toward the pial surface was visible. Since no information about ocularity is expected from extrastriate cortex, the exploited fMRI signal also needs to contain other information that enables classification. V2 and V3 were split in half to examine the depedendency on the distance to V1. Indeed, ***Figure 8*** shows a gradual performance decrease with larger distances to V1. This could be a hint to remaining partial volume contributions with V1 voxels due to the convoluted nature of the cerebral cortex.

**Figure 8.**
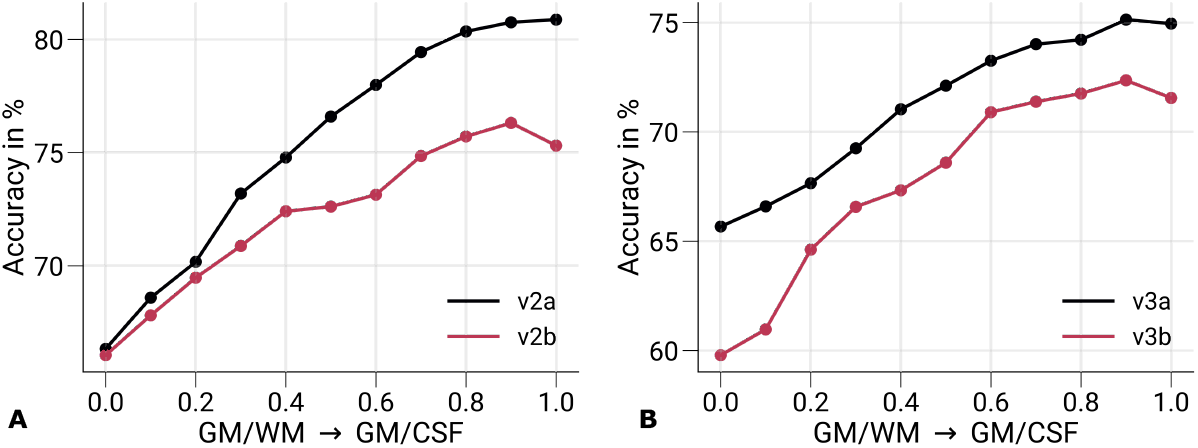
Prediction accuracies in V2 and V3. Mean prediction accuracies (prediction of the stimulated eye) for GE-BOLD are shown for V2 **(A)** and V3 **(B)** across cortical depth, respectively. Both areas were split in half based on retinotopy, with V2a and V3a being the half closer to V1. In **A–B**, data were significantly different (*p* < 0.05) from a 50% chance level at each cortical depth. The *p*-value was determined by bootstrapping (*n* = 1, 000) and corrected for multiple comparisons of individual layers (FDR correction using the Benjamini and Hochberg procedure). Decoding performance in areas V2 and V3 cannot be attributed to responses at the columnar level and indicate that also decoding performance in V1 may not be exclusively caused by responses at the columnar level. V2: secondary visual cortex, V3: tertiary visual cortex.

To exclude this alternative explanation, we ran an additional analysis, which is illustrated in ***Figure 9***. In brief, we computed the Euclidean distances between each vertex in V3 to its nearest vertex in V1 on the same surface for all participants. This was done both for GM/WM and GM/CSF surfaces, respectively. ***Figure 9*** shows that partial volume effects are unlikely to contribute to decoding accuracies from V3 regarding the used nominal voxel sizes used in fMRI acquisitions. However, it should be kept in mind that signal contributions might still leak into data sampled from neighboring areas due to the large physiological point-spread function of the BOLD signal (Engel, Glover, and Wandell, 1997; Parkes et al., 2005; Shmuel et al., 2007), which should be addressed in further studies.

**Figure 9.**
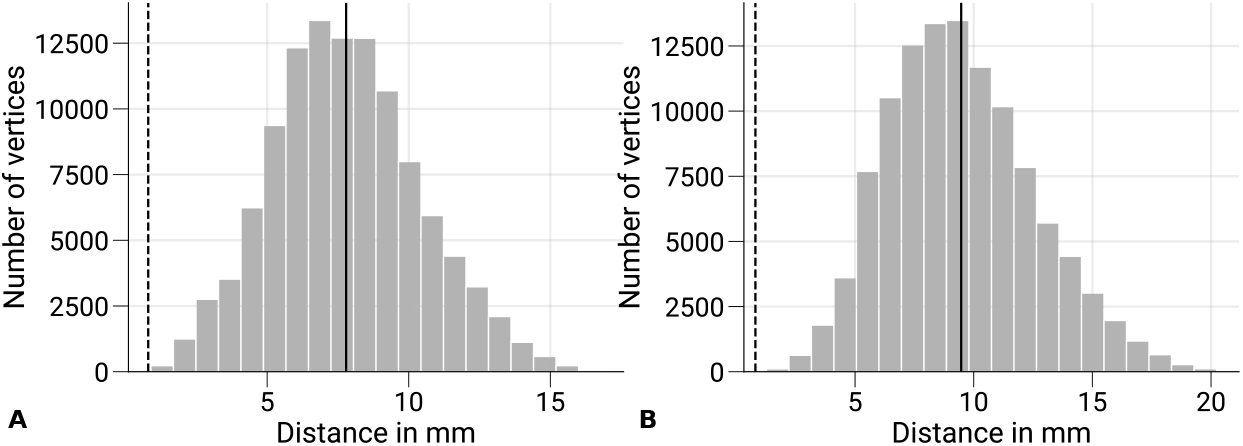
Minimum distances between V1 and sampled V3 data. The distribution of Euclidean distances between V3 vertices of the GM/WM **(A)** and the GM/CSF **(B)** and the closest V1 vertex of the same surface is shown across subjects and hemispheres. The overall mean is denoted as black vertical line and the nominal voxel size (0.8 mm) of functional acquisition is shown as vertical dashed line for reference. Voxel data sampled on V3 surfaces show minimal overlap with V1 regarding the used voxel size.

In VASO measurements, we exploit a CBV-weighted contrast that has a different temporal evolution compared to the BOLD response (Buxton, Wong, and Frank, 1998; Silva, Koretsky, and Duyn, 2007). More specifically, the CBV response has no initial dip, a shorter time-to-peak after stimulus onset, no poststimulus undershoot after stimulus offset, and needs more time to return to baseline. However, for the univariate analysis and the repeatability analysis, we processed data from all acquisition types with the same canonical HRF. As a control, we also analyzed the VASO data with a modified HRF that more closely resembled the CBV response’s time evolution (data not shown), which only resulted in minor differences to the presented results. Note that we did not use an HRF model for the multivariate analysis, since analysis was based on the steady-state time points in pre-processed fMRI time series.

One limitation of the experimental setup was that the used stimulus differed in color and luminance between eyes that was not explicitly accounted for. This might have led to decodable information along the parvo- and magnocellular streams inside but also outside of V1 (Tootell and Nasr, 2017). For example, ***Supplementary Figure 1***–***Supplementary Figure 5*** illustrate ODC maps from single participants, which generally show higher responses for the left eye, irrespective of eye dominance of single participants (eye dominance is stated in corresponding figure captions), which might be caused by remaining luminance differences between colors and therefore between eyes. Similar observations were made in an early fMRI decoding study, in which the eye-of-origin was decoded from a binocular rivalry stimulus (Haynes and Rees, 2005b). In binocular rivalry, the left and right eye receives incongruent stimuli, which were presented via anaglyph goggles. In that study, color filters were swapped between successive fMRI scanning runs in a control experiment. This resulted in decreased decoding performance in V1, whereas in extrastriate area V3 it stayed above chance level. From these results, it was concluded that performance in V1 was mostly based on ocularity information, while extrastriate areas V2 and V3 exploited more the color information in the stimulus. While not having the data to confirm these results in our experiment, we hypothesize that a similar effect contributed to the decodability in extrastriate areas as seen in ***Figure 8***.

Another limitation in the analysis is that data was pooled irrespective of visual field location. ODCs are known to vary in size and strength at different visual field locations (Adams, Sincich, and Horton, 2007), which might have influenced the results to some degree.

The acquired fMRI signal might therefore be influenced by several biases that are not related to ocularity information. These biases will also lead to differences in the expected laminar profile. However, we emphasize that, compared to other decoding studies exploiting information encoded at the columnar level with a conventional resolution, we could map and visualize ODCs in all single participants. That means that fine-grained information at the spatial scale of ODCs was present and the dominant pattern in univariate activation maps (see ***Figure 2***), which potentially could have been exploited by the linear classifier.

Our study analyzed the laminar specificity of MVPA with GE-BOLD, SE-BOLD, and VASO for the retrieval of information encoded at the spatial scale of cortical columns. For the first time, we used VASO in combination with MVPA to retrieve information from fine-grained cortical structures at the level of cortical layers. GE-BOLD is a very time-efficient acquisition method with larger SNR compared to SE-BOLD and VASO. This enables GE-BOLD to decode columnar information with high accuracy. However, the signal is weighted toward macrovascular signal contributions, limiting its capabilities to resolve information at the level of cortical layers. In comparison, VASO encodes two volumes at two inversion times, which limits its time efficiency. In addition, the BOLD correction in VASO is performed by a division operation, which enhances noise in the time series. This manifested itself in overall lower decoding accuracies for VASO.

In this regard, it might be a viable alternative to exploit the high SNR of GE-BOLD in combination with post-processing techniques to enhance the spatial specificity of the signal. Over the years, several approaches have been suggested that included deconvolution of cortical profiles (Markuerkiaga, Barth, and Norris, 2016; Hollander et al., 2021; Marquardt et al., 2020), masking out veins (Shmuel et al., 2007; Koopmans, Barth, and Norris, 2010; Moerel et al., 2018; Kay et al., 2019), spatial filtering of lower spatial frequencies of no interest (Sengupta et al., 2017; Mandelkow, Zwart, and Duyn, 2017; Hollander et al., 2021; Schmidt et al., 2024) or exploiting temporal information in the hemodynamic response (Kay et al., 2020) to remove macrovascular biases from GE-BOLD data. An extensive comparison between to these postprocessing steps is out of scope of the current study but might be an alternative route for decoding information at the mesoscopic scale based on acquisition techniques relying on the BOLD contrast.

In conclusion, the similar decoding profiles between acquisition techniques suggest that macroscopic venous effects are the predominant contributor that is exploited by the classifier in all cases. However, an exploratory analysis showed enhanced laminar specificity when using MVPA with VASO if the influence of feature selection is carefully considered. Future work is needed to further examine the potential increase in laminar specificity when combining multivariate techniques as MVPA with VASO.

## Acknowledgments

The research leading to these results has received funding from the European Research Council under the European Union’s Seventh Framework Program (FP7/2007-2013) /ERC grant agreement n° 616905. Nikolaus Weiskopf has received funding from the European Union’s Horizon 2020 research and innovation program under the grant agreement n° 681094 and from the BMBF (01EW1711A & B) in the framework of ERA-NET NEURON. Shahin Nasr has received funding from the NIH National Eye Institute (NEI) under the grant agreement n° R01EY030434. We thank the University of Minnesota Center for Magnetic Resonance Research for the provision of the multiband EPI sequence software. We thank Roland Mueller for the help with building the anaglyph spectacles. Furthermore, we thank Laurentius Huber for the provision of the SS-SI VASO sequence.

## Author contributions

Daniel Haenelt, Conceptualization, Methodology, Software, Formal analysis, Investigation, Data curation, Writing - original draft preparation, Writing - review & editing, Visualization; Denis Chaimow, Conceptualization, Methodology, Writing - review & editing; Marianna Elisa Schmidt, Formal analysis, Writing - review & editing; Shahin Nasr, Methodology, Software, Writing - review & editing; Nikolaus Weiskopf, Conceptualization, Resources, Writing - review & editing, Supervision, Project administration, Funding acquisition; Robert Trampel, Investigation, Writing - review & editing, Supervision

## Supplementary Information

**Supplementary Figure 1.**
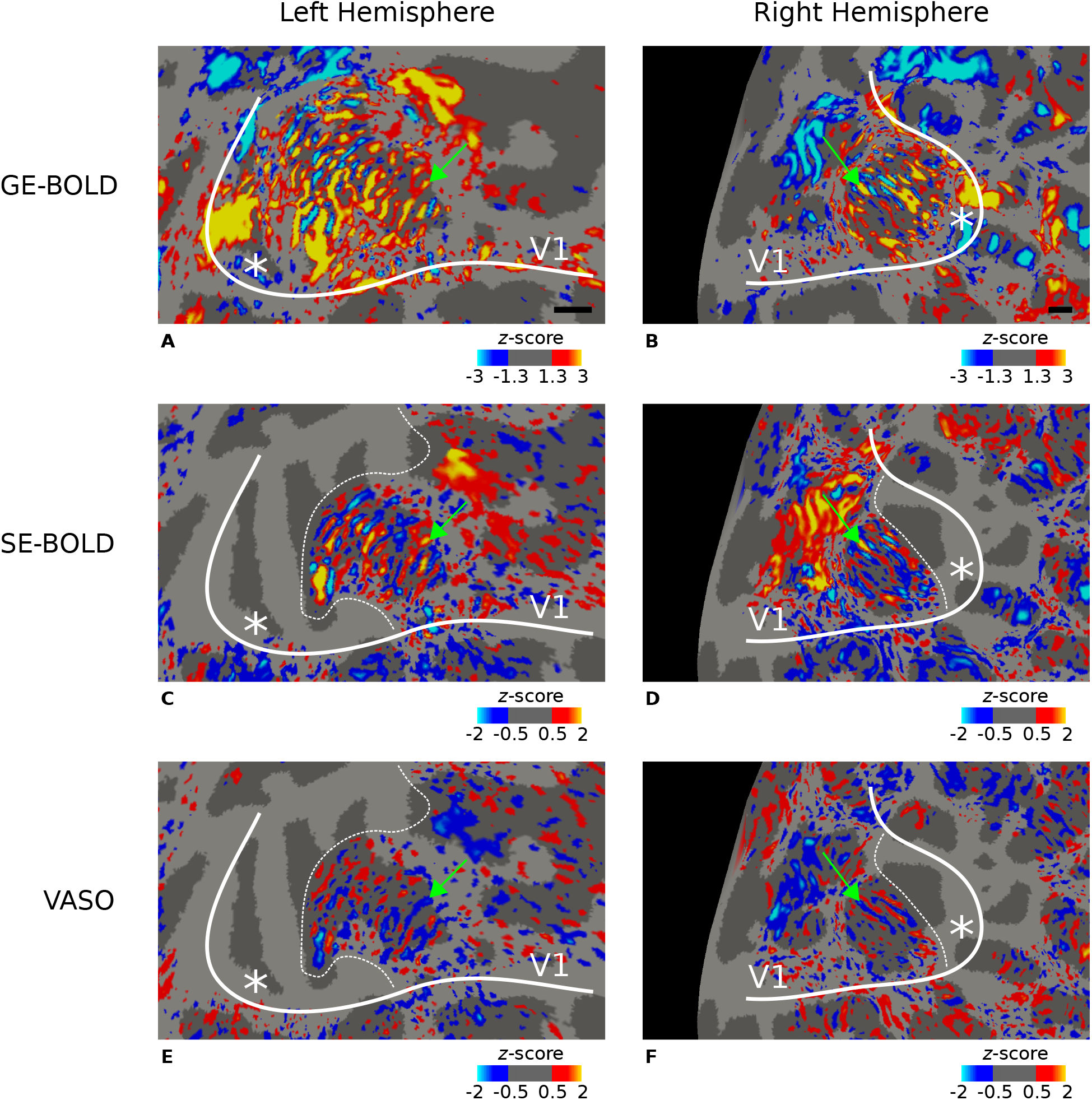
Ocular dominance columns (ODCs) from subject 1. Thresholded activation maps (contrast left eye > right eye) are shown for the left and right hemisphere, respectively, for GE-BOLD **(A–B)**, SE-BOLD **(C–D)**, and VASO **(E–F)**. Data were averaged across sessions, sampled at mid-cortical depth, and shown on flattened surfaces. Similarities between maps are evident. Green arrows point to columns that were reproducibly activated between scanning sessions. This participant was left eye dominant. Note that VASO has an inverted contrast compared to BOLD. Black lines in **A** and **B** show scale bars (5 mm), respectively—other details as in ***Figure 2***.

**Supplementary Figure 2.**
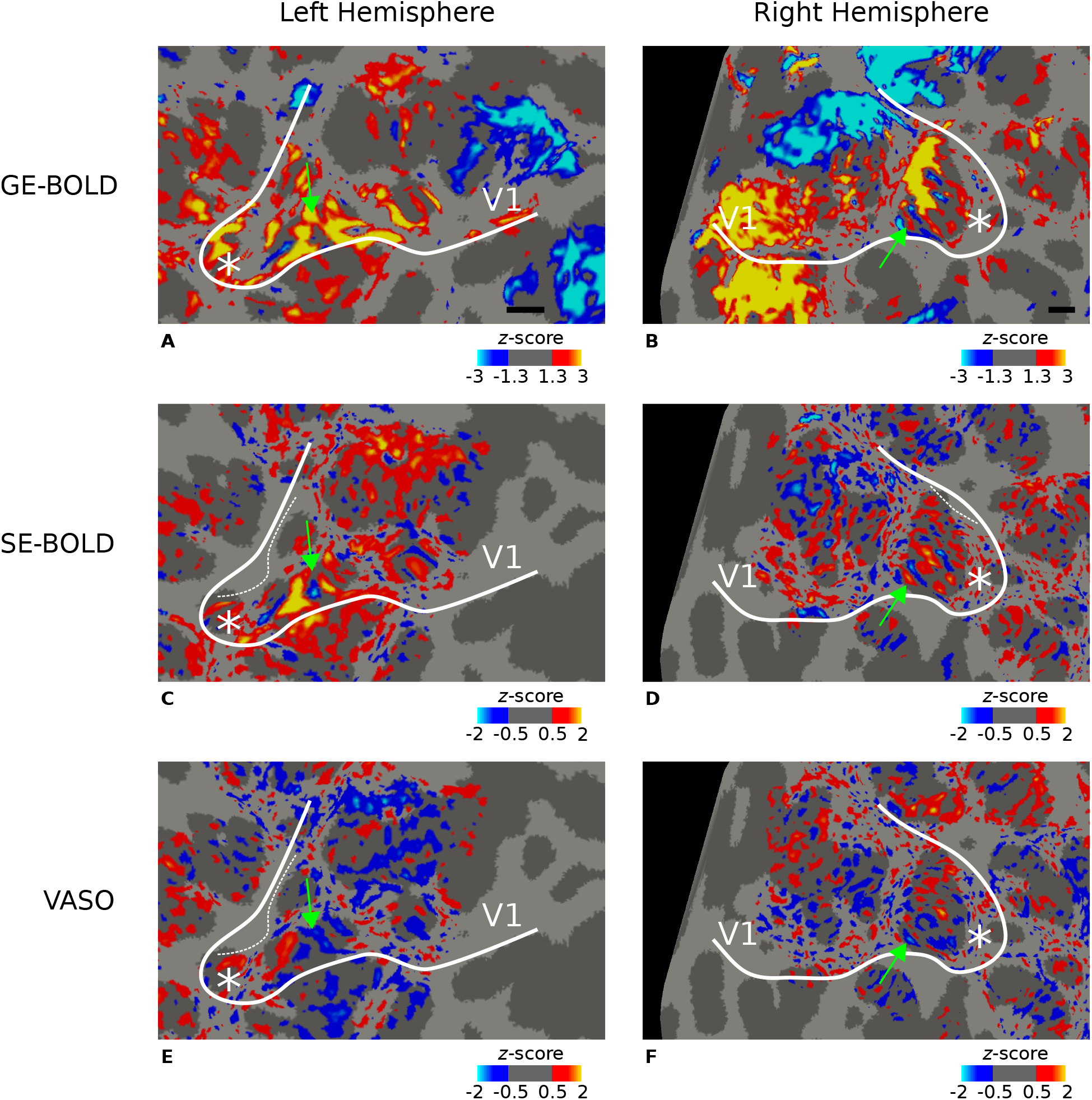
Ocular dominance columns (ODCs) from subject 2. Thresholded activation maps (contrast left eye > right eye) are shown for the left and right hemisphere, respectively, for GE-BOLD **(A–B)**, SE-BOLD **(C–D)**, and VASO **(E–F)**. Data were averaged across sessions, sampled at mid-cortical depth, and shown on flattened surfaces. Similarities between maps are evident. Green arrows point to columns that were reproducibly activated between scanning sessions. This participant was right eye dominant. Note that VASO has an inverted contrast compared to BOLD. Black lines in **A** and **B** show scale bars (5 mm), respectively—other details as in ***Figure 2***.

**Supplementary Figure 3.**
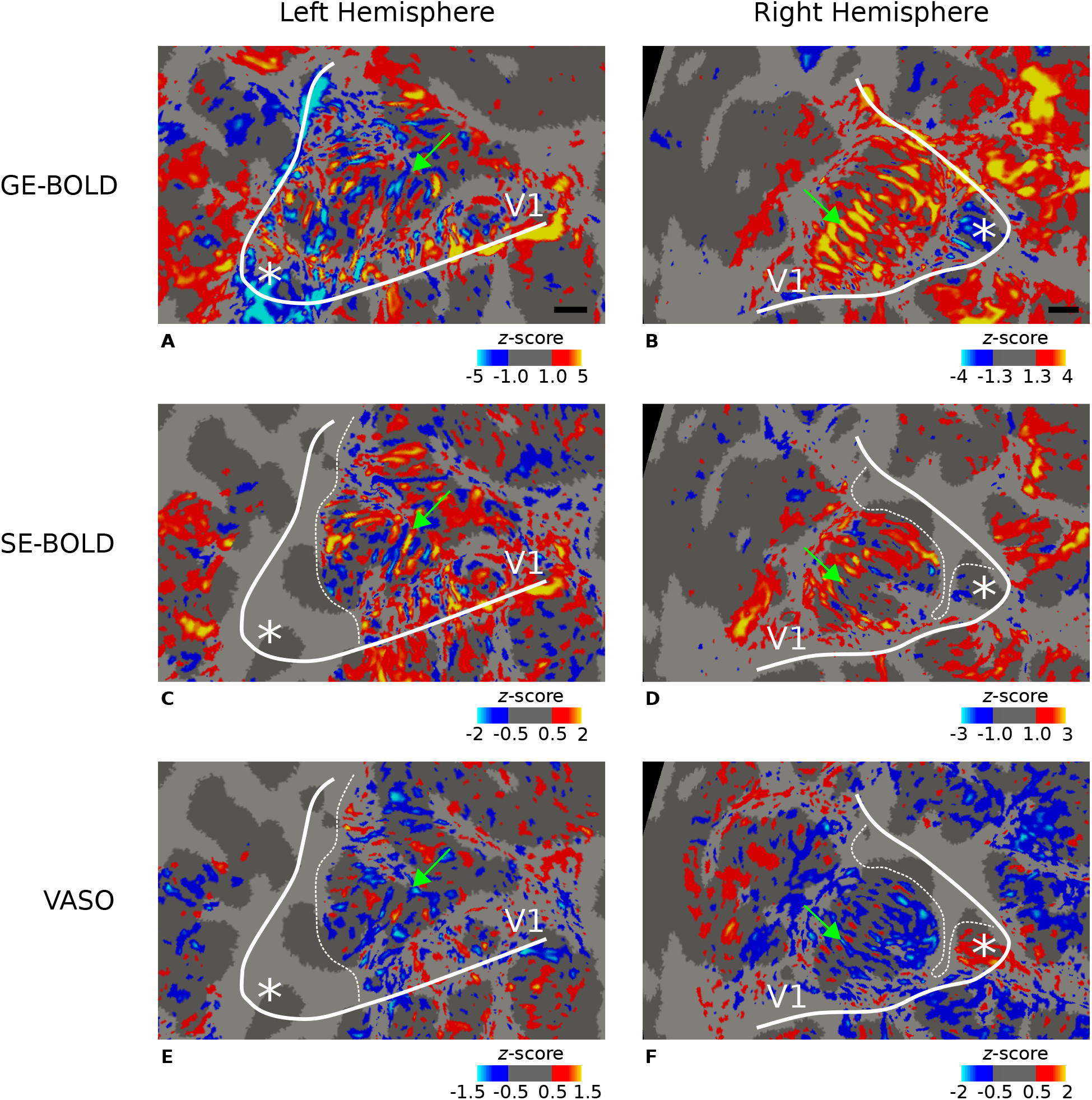
Ocular dominance columns (ODCs) from subject 3. Thresholded activation maps (contrast left eye > right eye) are shown for the left and right hemisphere, respectively, for GE-BOLD **(A–B)**, SE-BOLD **(C–D)**, and VASO **(E–F)**. Data were averaged across sessions, sampled at mid-cortical depth, and shown on flattened surfaces. Similarities between maps are evident. Green arrows point to columns that were reproducibly activated between scanning sessions. This participant was left eye dominant. Note that VASO has an inverted contrast compared to BOLD. Black lines in **A** and **B** show scale bars (5 mm), respectively—other details as in ***Figure 2***.

**Supplementary Figure 4.**
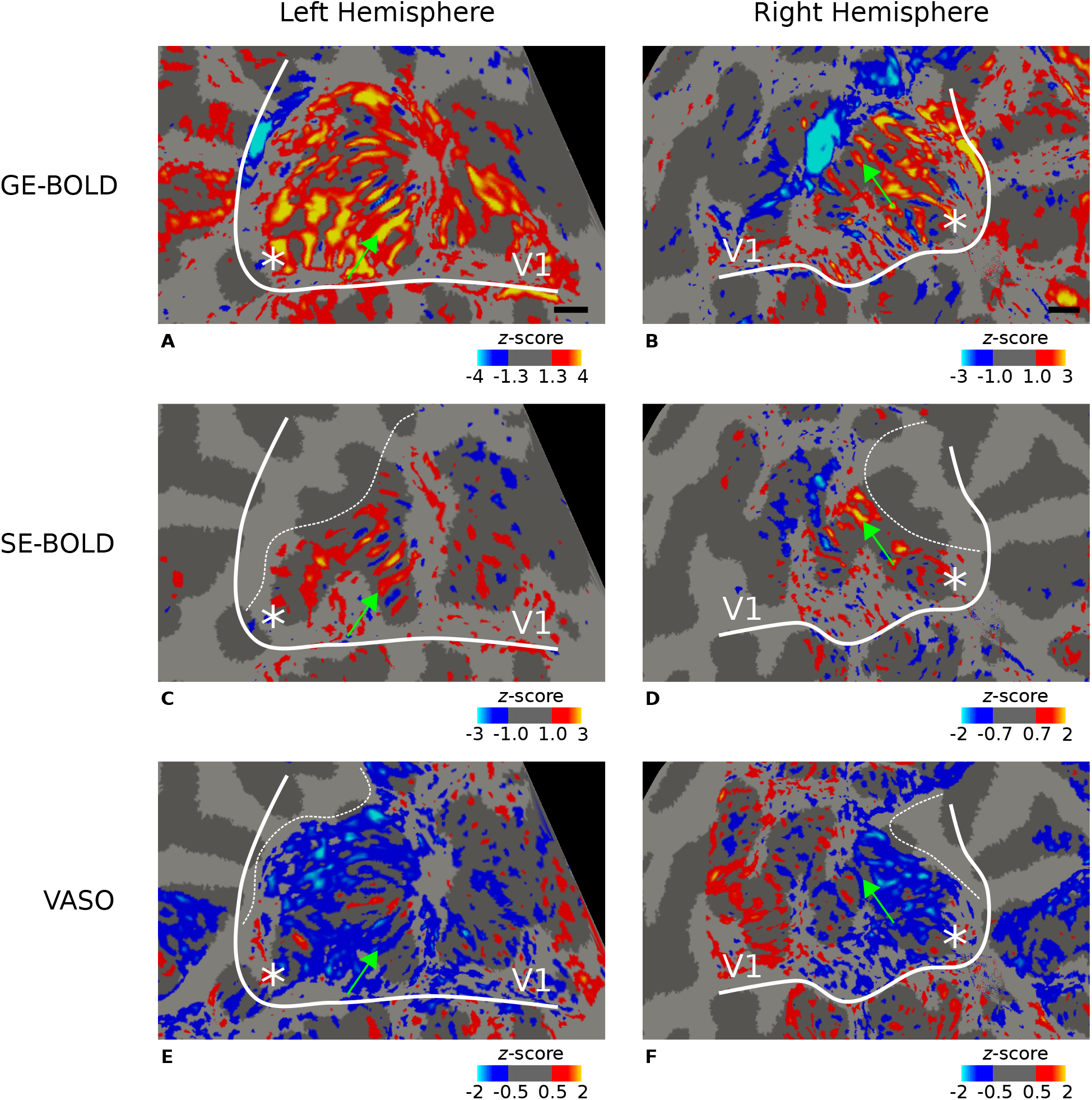
Ocular dominance columns (ODCs) from subject 4. Thresholded activation maps (contrast left eye > right eye) are shown for the left and right hemisphere, respectively, for GE-BOLD **(A–B)**, SE-BOLD **(C–D)**, and VASO **(E–F)**. Data were averaged across sessions, sampled at mid-cortical depth, and shown on flattened surfaces. Similarities between maps are evident. Green arrows point to columns that were reproducibly activated between scanning sessions. This participant was right eye dominant. Note that VASO has an inverted contrast compared to BOLD. Black lines in **A** and **B** show scale bars (5 mm), respectively—other details as in ***Figure 2***.

**Supplementary Figure 5.**
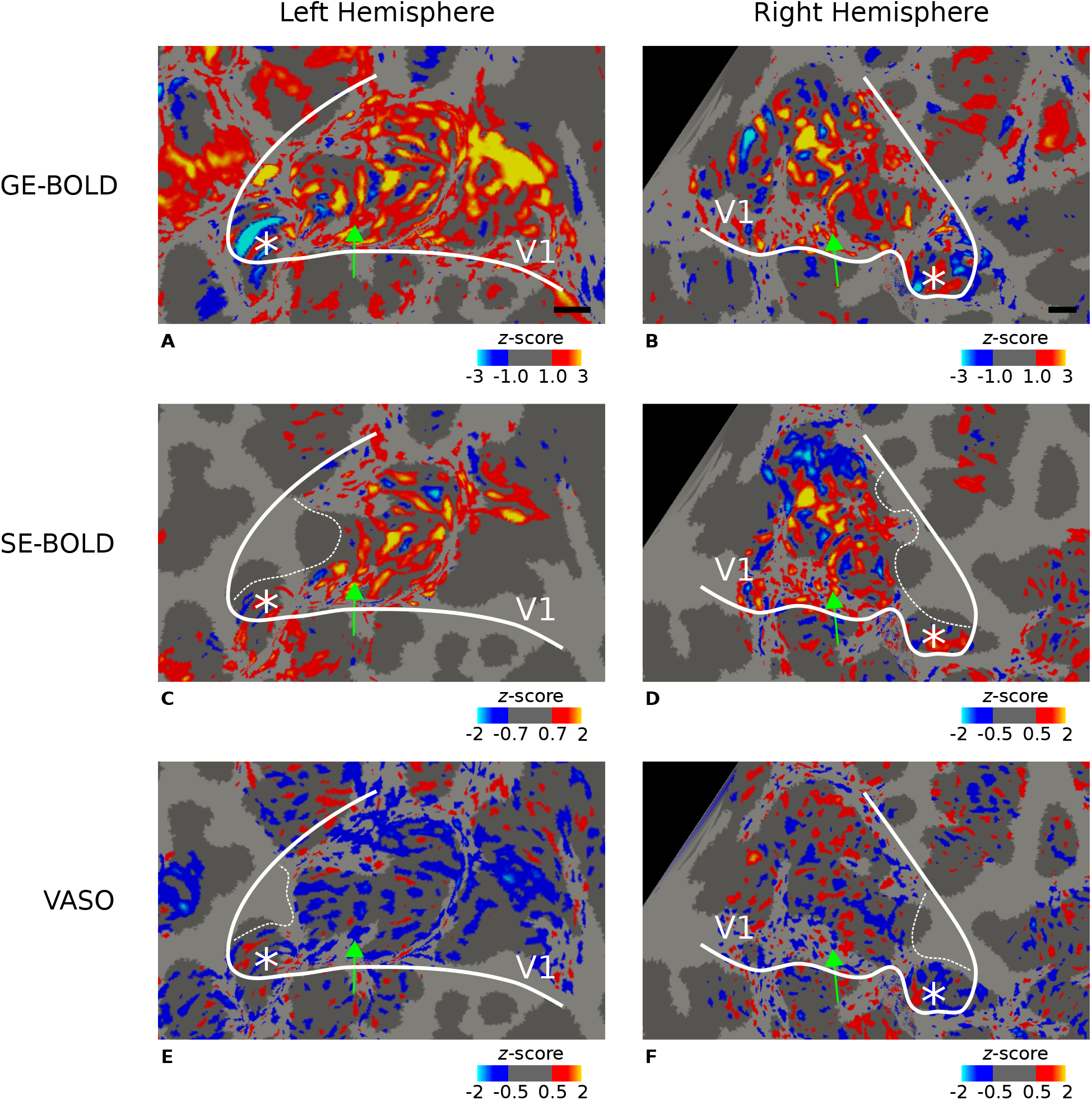
Ocular dominance columns (ODCs) from subject 5. Thresholded activation maps (contrast left eye > right eye) are shown for the left and right hemisphere, respectively, for GE-BOLD **(A–B)**, SE-BOLD **(C–D)**, and VASO **(E–F)**. Data were averaged across sessions, sampled at mid-cortical depth, and shown on flattened surfaces. Similarities between maps are evident. Green arrows point to columns that were reproducibly activated between scanning sessions. This participant was left eye dominant. Note that VASO has an inverted contrast compared to BOLD. Black lines in **A** and **B** show scale bars (5 mm), respectively—other details as in ***Figure 2***.

**Supplementary Figure 6.**
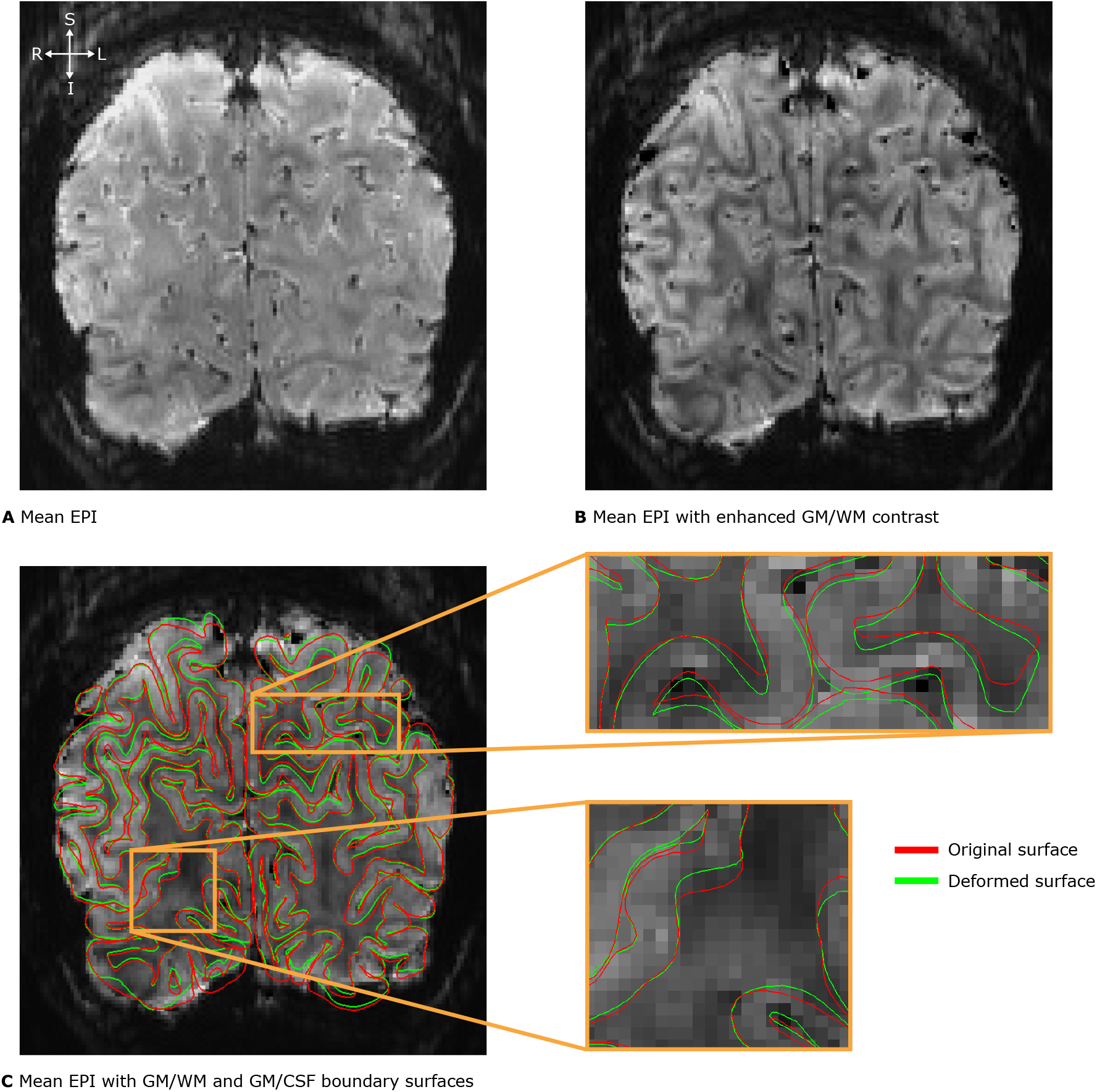
Illustration of the GBB method. The method is used to enhance the alignment of cortical boundary surfaces based on an undistorted whole-brain anatomy to the cortical borders found in distorted functional images. **A** shows the temporal mean of the functional time series without task (GE-BOLD, 200 time points, subject 3) in coronal view that was acquired in the first session. **B** To enhance the GM/WM border and thereby increase the robustness of the proposed method, we weighted the temporal mean by its phase (see to Data preprocessing for detailed information) as usually done in susceptibility-weighted imaging methods. In **C**, the surfaces before (depicted in red) and after (depicted in green) alignment with the GBB method are presented. This technique is implemented in the GBB package (0.1.6, https://pypi.org/project/gbb/). The core idea of the method is to locally deform the GM/WM boundary surface iteratively until it reaches the GM/WM border found in the functional data. Each iteration starts by randomly selecting one vertex. Then, the vertex and its surrounding neighborhood is moved a small amount along the direction of increased GM/WM contrast scaled by a set step size. The change is evaluated by using the same cost function proposed in Greve and Fischl, 2009. Before alignment, surfaces are transformed into functional space via a rigid registration. From resulting vertex displacements of the GM/WM border, a deformation field is estimated that is then applied to the GM/CSF surface. The method improves spatial correspondence between the surfaces and the GM/WM boundaries observed in the functional images. GM: gray matter, WM: white matter, CSF: cerebrospinal fluid.

**Supplementary Figure 7.**
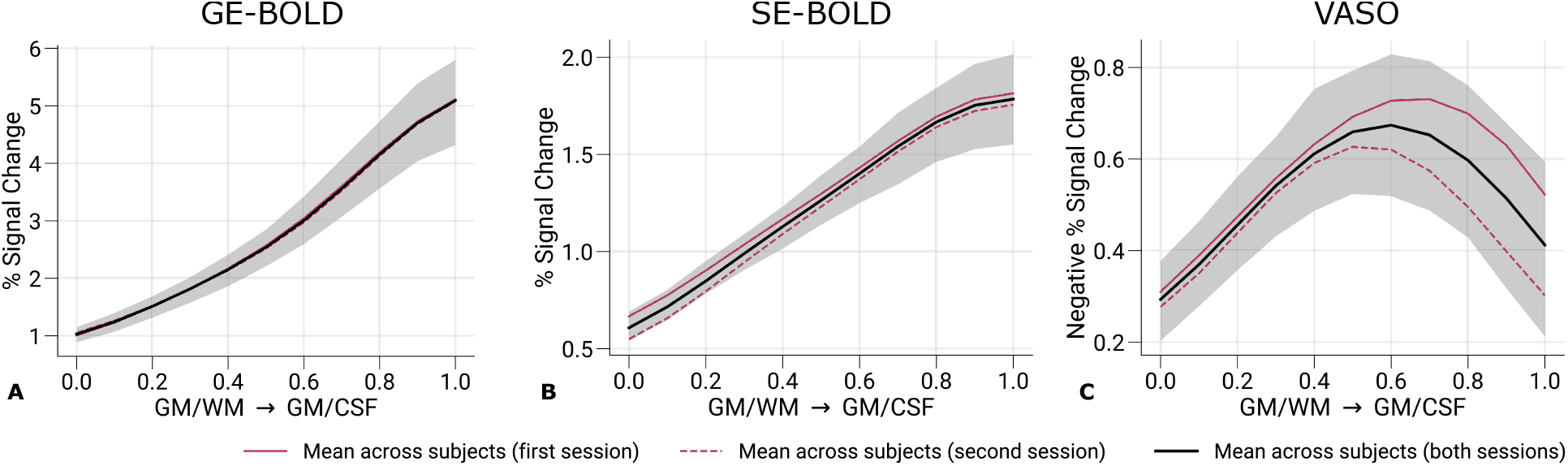
Percent signal changes across cortical depth from whole V1. Mean percent signal changes (contrast: left eye and right eye > baseline) for GE-BOLD **(A)**, SE-BOLD **(B)**, and VASO **(C)** are shown across cortical depth. Contrary to ***Figure 5***, all V1 data inside the field of view across all scanning sessions were used. Compared to ***Figure 5***, lower percent signal changes and lower variability across participants can be identified. In **C**, the peak at mid-cortical depth is more pronounced. Note that we inverted the y-axis in **C** for consistency with **A** and **B**—other details as in ***Figure 5***.

**Supplementary Figure 8.**
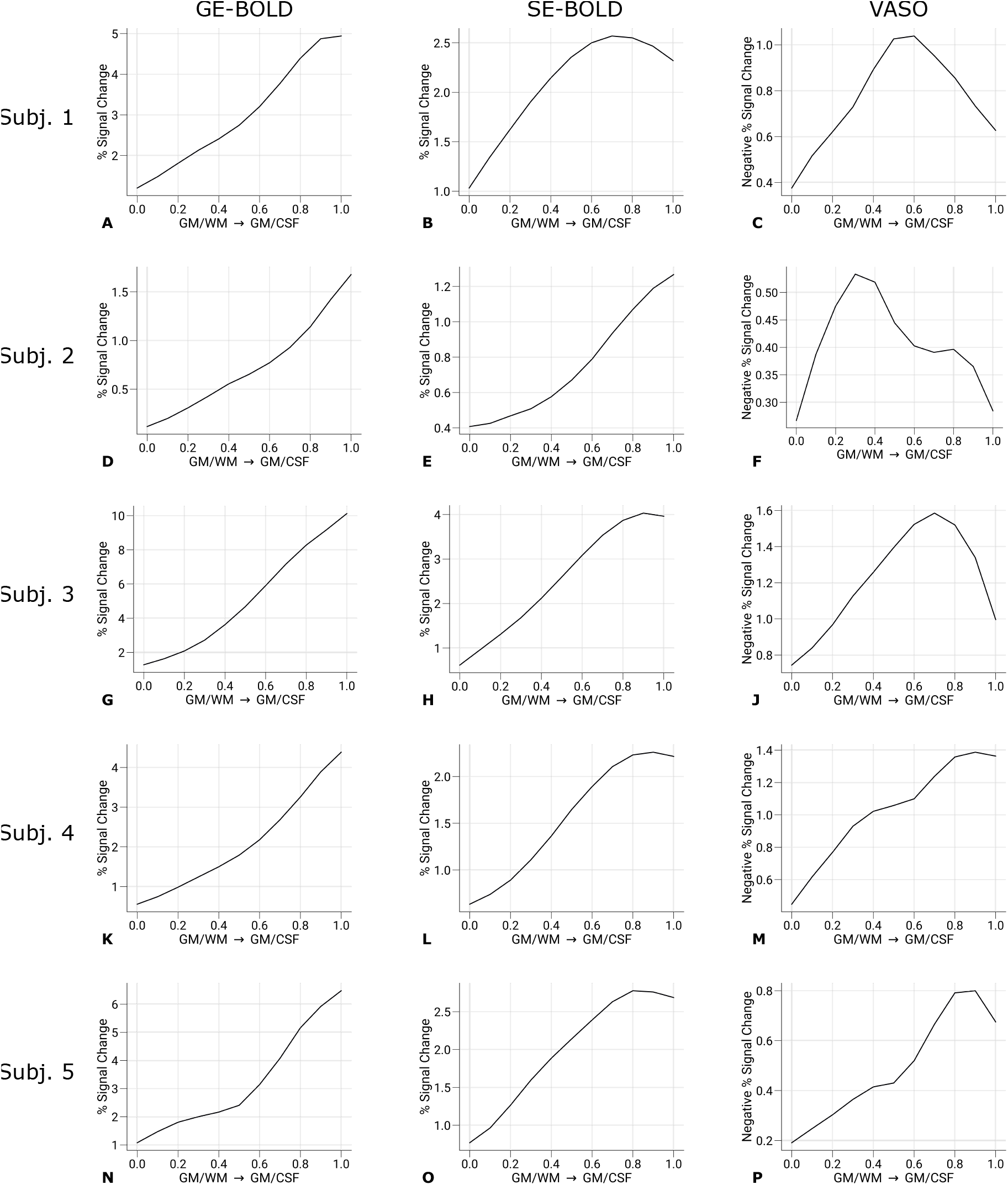
Percent signal changes across cortical depth from single participants. Percent signal changes (contrast: left eye and right eye > baseline) for GE-BOLD (left column), SE-BOLD (middle column), and VASO (right column) are shown across cortical depth for single participants (average across two sessions). Only data points (n = 200) were used that were also selected for the decoding analysis. Note that we inverted the y-axis for VASO (right column) for easier interpretation. The variability of cortical profiles between participants can be identified.

**Supplementary Figure 9.**
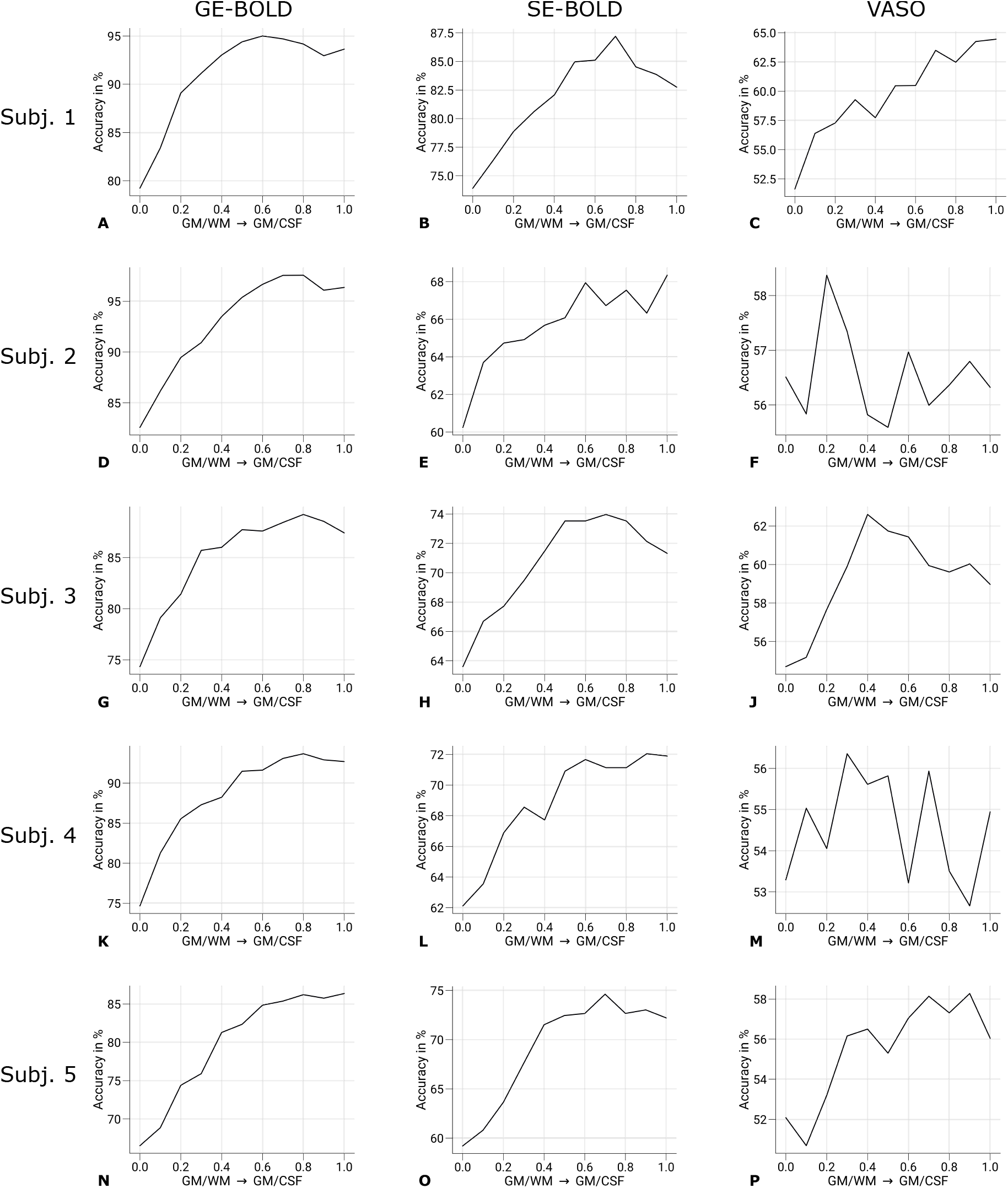
Prediction accuracies across cortical depth from single participants. Prediction accuracies (prediction of the stimulated eye) for GE-BOLD (left column), SE-BOLD (middle column), and VASO (column) are shown across cortical depth for single participants (average across two sessions). The variability of cortical profiles between participants can be identified.

**Supplementary Figure 10.**
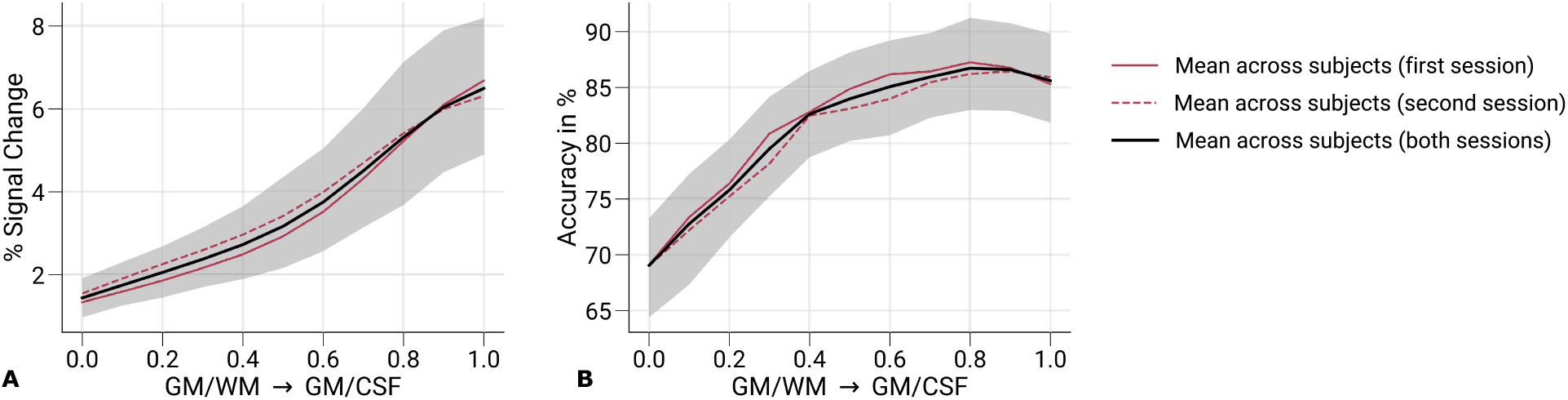
Percent signal changes and prediction accuracies for not-nulled time points in VASO sessions. Mean percent signal changes (contrast: left eye and right eye > baseline) **(A)** and mean prediction accuracies (prediction of the stimulated eye) **(B)** are shown across cortical depth for not-nulled (BOLD-weighted) time series from VASO sessions. Red solid and dashed lines show the mean across participants from the first and second session, respectively. Black lines indicate the mean across participants and scanning sessions. The gray area demarcates the bootstrap 95% confidence interval (n = 1,000). Shapes of cortical profiles are similar to ***Figure 5A*** and ***Figure 6A***, respectively. Overall, lower prediction accuracies compared to ***Figure 6A*** might be attributable to the smaller temporal efficiency due to the longer TR in VASO acquisitions.

**Supplementary Figure 11.**
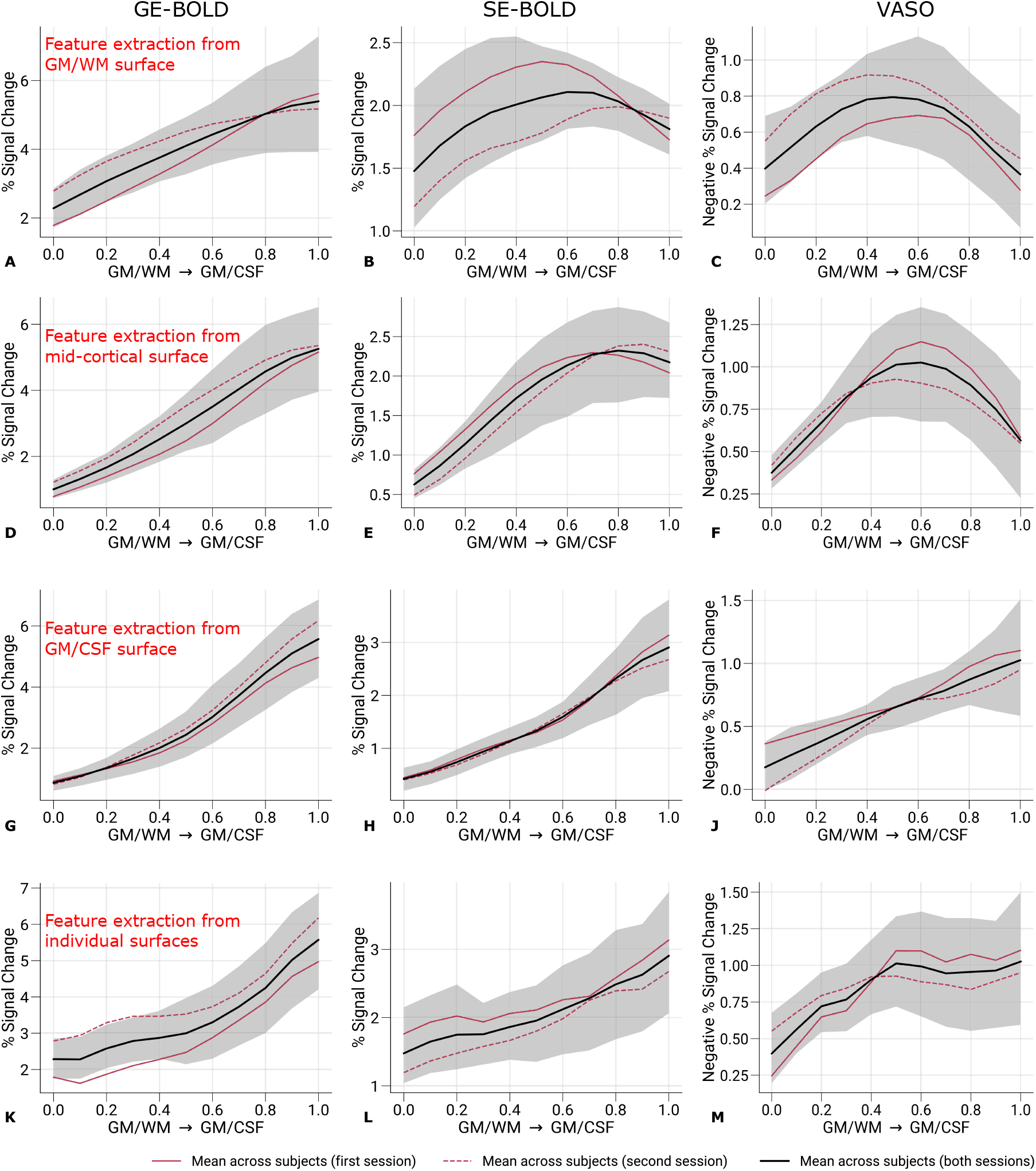
Percent signal changes across cortical depth. Mean percent signal changes (contrast: left eye and right eye > baseline) for GE-BOLD (left column), SE-BOLD (middle column), and VASO (right column) are shown across cortical depth. In contrast to ***Figure 5***, features selection was restricted to data points sampled on the GM/WM **(A–C)**, the mid-cortical **(D–F)**, and the GM/CSF **(G–J)** boundary surfaces, respectively. In **K–M**, feature selection was performed for each cortical layer independently—other details as in ***Figure 5***.

**Supplementary Figure 12.**
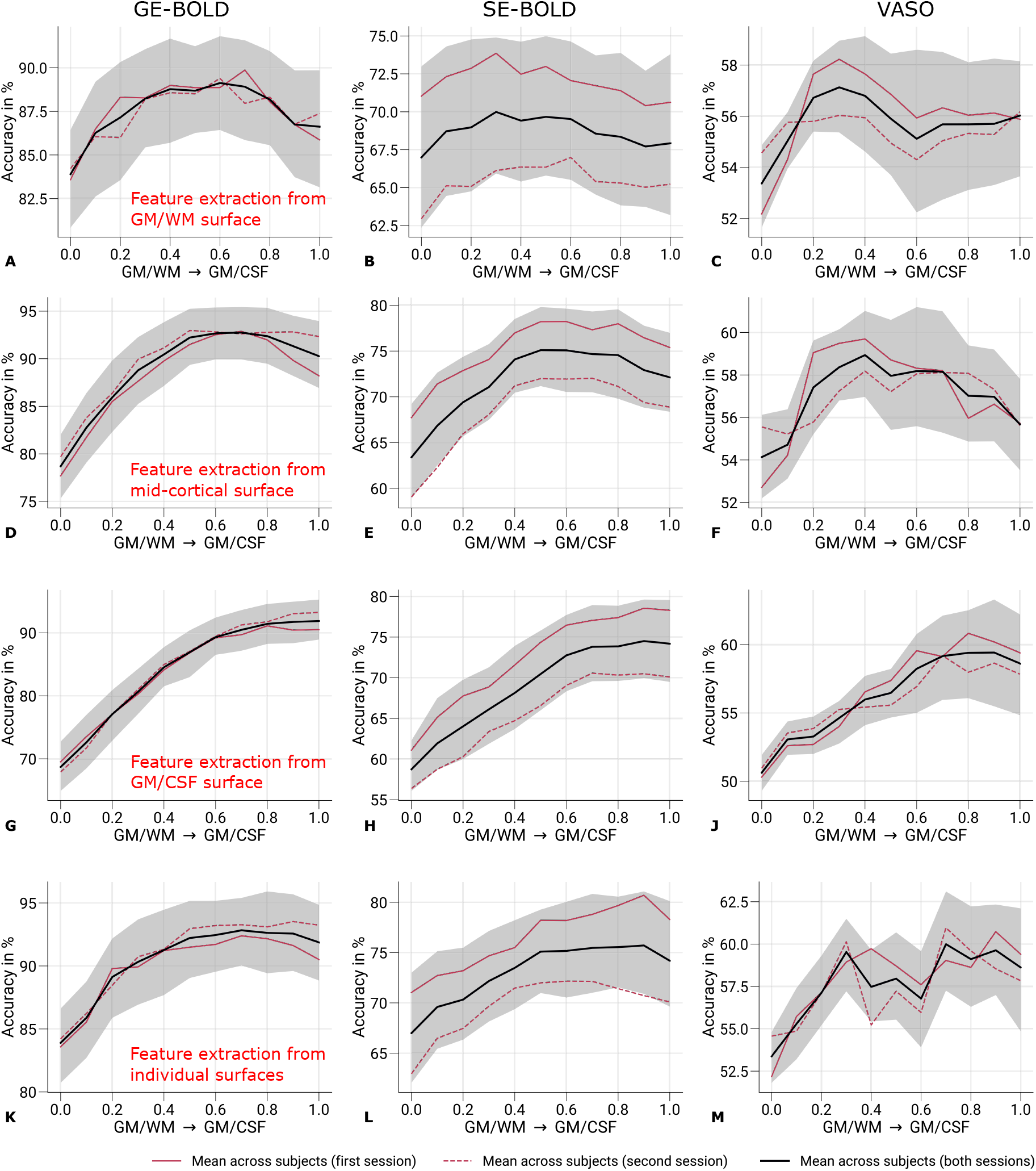
Prediction accuracies across cortical depth. Mean prediction accuracies (prediction of the stimulated eye) for GE-BOLD (left column), SE-BOLD (middle column), and VASO (right column) are shown across cortical depth. In contrast to ***Figure 6***, features selection was restricted to data points sampled on the GM/WM **(A–C)**, the mid-cortical **(D–F)**, and the GM/CSF **(G–J)** boundary surfaces, respectively. In **K–M**, feature selection was performed for each cortical layer independently—other details as in ***Figure 6***.

**Supplementary Figure 13.**
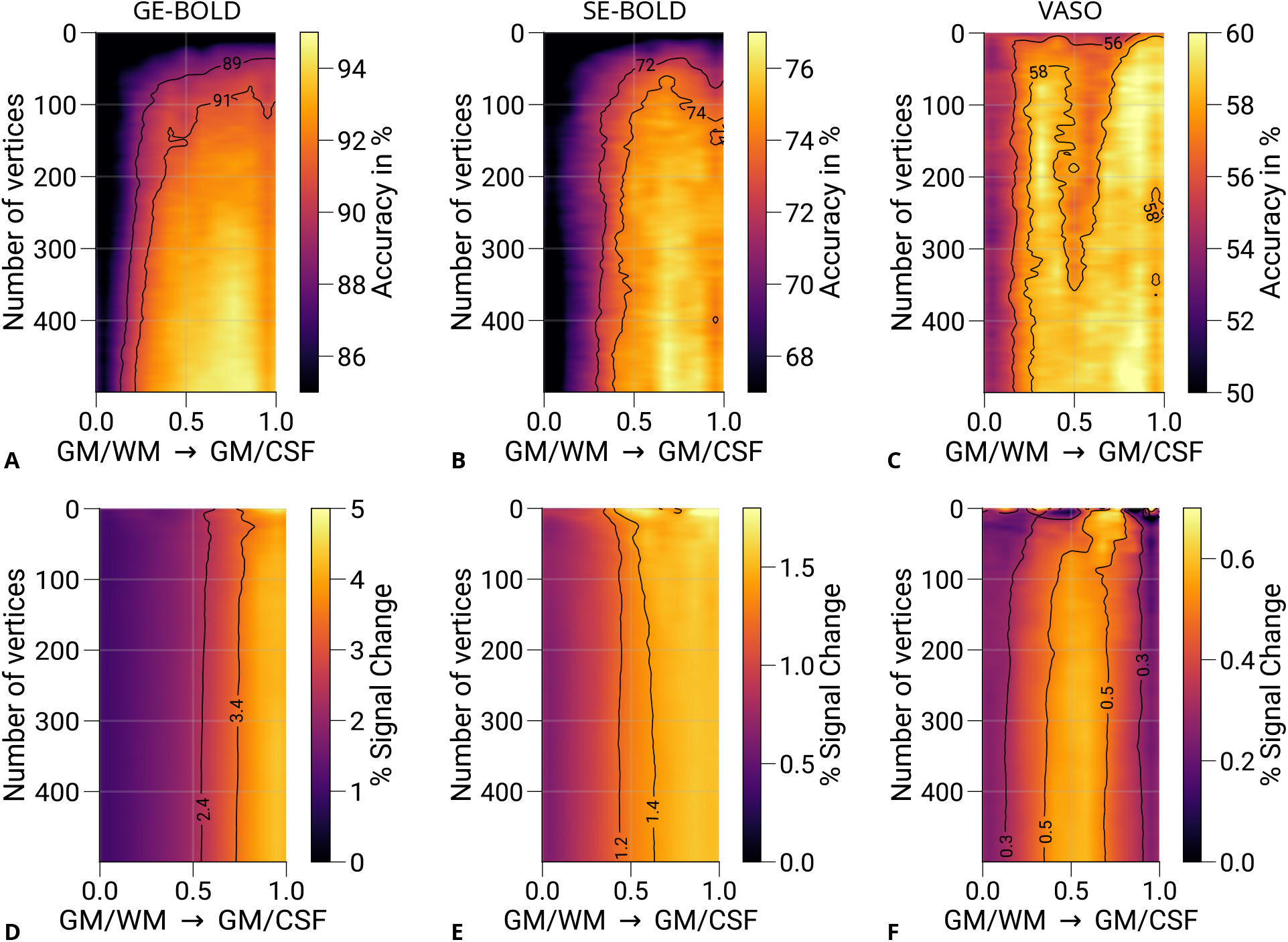
Prediction accuracies and percent signal changes for different number of features. Mean prediction accuracies (prediction of the stimulated eye) for GE-BOLD **(A)**, SE-BOLD **(B)**, and VASO **(C)** are shown for a varying number of features (vertices) across cortical depth. **D–F** show corresponding percent signal changes (left eye and right eye > baseline) using the same data points. In contrast to ***Figure 7***, feature selection was performed for each cortical layer independently—other details as in ***Figure 7***.

## Footnotes

1 For the decoding analysis, highpass filtering was based on an in-house filter that convolved the time series with a Gaussian running line smoother. For all other analyses, highpass filtering was performed with SPM12.

2 Note that the phase data was solely used for moving the boundary surfaces during preprocessing and was not considered in further analyses.

3 Since the cortical surfaces were defined in a spatially distorted fMRI space, the equidistant approach was preferred over the more anatomically precise equivolume approach (Waehnert et al., 2014).

## References

Abdul-Rahman, Hussein, Munther Gdeisat, David Burton, and Michael Lalor (2005). “Fast three-dimensional phase-unwrapping algorithm based on sorting by reliability following a non-continuous path”. In: Optical Measurement Systems for Industrial Inspection IV. Ed. by Wolfgang Osten, Christophe Gorecki, and Erik L. Novak. Vol. 5856. International Society for Optics and Photonics. SPIE, pp. 32–40. DOI: 10.1117/12.611415.

Adams, Daniel L. and Jonathan C. Horton (2009). “Ocular dominance columns: enigmas and challenges”. In: The Neuroscientist 15.1, pp. 62–77. DOI: 10.1177/1073858408327806.

Adams, Daniel L., Lawrence C. Sincich, and Jonathan C. Horton (2007). “Complete pattern of ocular dominance columns in human primary visual cortex”. In: Journal of Neuroscience 27.39, pp. 10391–10403. DOI: 10.1523/JNEUROSCI.2923-07.2007.

Akbari, Atena, Joseph S. Gati, Peter Zeman, Brett Liem, and Ravi S. Menon (2023). “Layer dependence of monocular and binocular responses in human ocular dominance columns at 7T using VASO and BOLD”. In: bioRxiv. DOI: 10.1101/2023.04.06.535924. URL: https://www.biorxiv.org/content/10.1101/2023.04.06.535924.

Andersson, Jesper L. R., Stefan Skare, and John Asburner (2003). “How to correct susceptibility distortions in spin-echo echo-planar images: application to diffusion tensor imaging”. In: NeuroImage 20.2, pp. 870–888. DOI: 10.1016/S1053-8119(03)00336-7.

Andrews, Timothy J., Scott D. Halpern, and Dale Purves (1997). “Correlated size variations in human visual cortex, lateral geniculate nucleus, and optic tract”. In: Journal of Neuroscience 17.8, pp. 2859–2868. DOI: 10.1523/JNEUROSCI.17-08-02859.1997.

Avants, Brian B., Charles L. Epstein, Murray Grossman, and James C. Gee (2008). “Symmetric diffeomorphic image registration with cross-correlation: evaluating automated labeling of elderly and neurodegenerative brain”. In: Medical Image Analysis 12.1, pp. 26–41. DOI: 10.1016/j.media.2007.06.004.

Beeck, Hans P. Op de (2010). “Against hyperacuity in brain reading: spatial smoothing does not hurt multivariate fMRI analyses?” In: NeuroImage 49.3, pp. 1943–1948. DOI: 10.1016/j.neuroimage.2009.02.047.

Boxerman, Jerrold L., Leena M. Hamberg, Bruce R. Rosen, and Robert M. Weisskoff (1995). “MR contrast due to intravascular magnetic susceptibility perturbations”. In: Magnetic Resonance in Medicine 34.4, pp. 555–566. DOI: 10.1002/mrm.1910340412.

Boynton, Geoffrey M. (2005). “Imaging orientation selectivity: decoding conscious perception in V1”. In: Nature Neuroscience 8.5, pp. 541–542. DOI: 10.1038/nn0505-541.

Brainard, David H. (1997). “The psychophysics toolbox”. In: Spatial Vision 10.4, pp. 433–436. DOI:10.1163/156856897X00357.

Brodmann, Korbinian (1909). Vergleichende Lokalisationslehre der Grosshirnrinde in ihren Prinzipien dargestellt auf Grund des Zellenbaues. Leipzig, Germany: J.A. Barth.

Burt, Joshua B., Markus Helmer, Maxwell Shinn, Alan Anticevic, and John D. Murray (2020). “Generative modeling of brain maps with spatial autocorrelation”. In: NeuroImage 220, p. 117038. DOI: 10.1016/j.neuroimage.2020.117038.

Buxton, Richard B. (2013). “The physics of functional magnetic resonance imaging (fMRI)”. In: Reports on Progress in Physics 76.9, p. 096601. DOI: 10.1088/0034-4885/76/9/096601.

Buxton, Richard B., Eric C. Wong, and Lawrence R. Frank (1998). “Dynamics of blood flow and oxygenation changes during brain activation: the balloon model”. In: Magnetic Resonance in Medicine 39.6, pp. 855–864. DOI: 10.1002/mrm.1910390602.

Chaimow, Denis, Essa Yacoub, Kâmil Uğurbil, and Amir Shmuel (2011). “Modeling and analysis of mechanisms underlying fMRI-based decoding of information conveyed in cortical columns”. In: NeuroImage 56.2, pp. 627–642. DOI: 10.1016/j.neuroimage.2010.09.037.

Chang, Chih-Chung and Chih-Jen Lin (2011). “LIBSVM: a library for support vector machines”. In: ACM Transactions on Intelligent Systems and Technology 2.3. DOI: 10.1145/1961189.1961199.

Cheng, Kang, R. Allen Waggoner, and Keiji Tanaka (2001). “Human ocular dominance columns as revealed by high-field functional magnetic resonance imaging”. In: Neuron 32.2, pp. 359–374. DOI: 10.1016/S0896-6273(01)00477-9.

Cox, Robert W. (1996). “AFNI: software for analysis and visualization of functional magnetic resonance neuroimages”. In: Computers and Biomedical Research 29.3, pp. 162–173. DOI: 10.1006/cbmr.1996.0014.

Dale, Anders M., Bruce Fischl, and Martin I. Sereno (1999). “Cortical surface-based analysis: I. segmentation and surface reconstruction”. In: NeuroImage 9.2, pp. 179–194. DOI: 10.1006/nimg.1998.0395.

Dechent, Peter and Jens Frahm (2000). “Direct mapping of ocular dominance columns in human primary visual cortex”. In: NeuroReport 11.14, pp. 3247–3249. DOI: 10.1097/00001756-200009280-00039.

Dobkins, Karen R., Alex Thiele, and Thomas D. Albright (2000). “Comparison of red–green equiluminance points in humans and macaques: evidence for different L:M cone ratios between species”. In: Journal of the Optical Society of America A 17.3, pp. 545–556. DOI: 10.1364/josaa.17.000545.

Dougherty, Kacie, Michele A. Cox, Jacob A. Westerberg, and Alexander Maier (2019). “Binocular modulation of monocular V1 neurons”. In: Current Biology 29.3, pp. 381–391. DOI: 10.1016/j.cub.2018.12.004.

Duvernoy, Henri M., S. Delon, and J. L. Vannson (1981). “Cortical blood vessels of the human brain”. In: Brain Research Bulletin 7.5, pp. 519–579. DOI: 10.1016/0361-9230(81)90007-1.

Engel, Stephen A., Gary H. Glover, and Brian A. Wandell (1997). “Retinotopic organization in human visual cortex and the spatial precision of functional MRI”. In: Cerebral Cortex 7.2, pp. 181–192. DOI: 10.1093/cercor/7.2.181.

Feinberg, David A., Steen Moeller, Stephen M. Smith, Edward Auerbach, Sudhir Ramanna, Matt F. Glasser, Karla L. Miller, Kâmil Uğurbil, and Essa Yacoub (2010). “Multiplexed echo planar imaging for sub-second whole brain fMRI and fast diffusion imaging”. In: PloS ONE 5.12, e15710. DOI: 10.1371/journal.pone.0015710.

Feinberg, David A., Salvatore Torrisi, Alexander J. S. Becket, Rüdiger Stirnberg, Tony Stöcker, Philipp Ehses, and Renzo Huber (2022). “Sub-0.1 microliter CBV fMRI on the Next Generation 7T scanner”. In: Proceedings International Society for Magnetic Resonance in Medicine. London, UK.

Feinberg, David A., An T. Vu, and Alexander Beckett (2018). “Pushing the limits of ultra-high resolution human brain imaging with SMS-EPI demonstrated for columnar level fMRI”. In: NeuroImage 164, pp. 155–163. DOI: 10.1016/j.neuroimage.2017.02.020.

Felleman, Daniel J. and David C. Van Essen (1991). “Distributed hierarchical processing in the primate cerebral cortex”. In: Cerebral Cortex 1.1, pp. 1–47. DOI: 10.1093/cercor/1.1.1-a.

Fischl, Bruce and Anders M. Dale (2000). “Measuring the thickness of the human cerebral cortex from magnetic resonance images”. In: Proceedings of the National Academy of Sciences 97.20, pp. 11050–11055. DOI: 10.1073/pnas.200033797.

Fischl, Bruce, Martin I. Sereno, and Anders M. Dale (1999). “Cortical surface-based analysis. II: inflation, flattening, and a surface-based coordinate system”. In: NeuroImage 9.2, pp. 195–207. DOI: 10.1006/nimg.1998.0396.

Formisano, Elia and Nikolaus Kriegeskorte (2012). “Seeing patterns through the hemodynamic veil — the future of pattern-information fMRI”. In: NeuroImage 62.2, pp. 1249–1256. DOI: 10.1016/j.neuroimage.2012.02.078.

Fracasso, Alessio, Natalia Petridou, and Serge O. Dumoulin (2016). “Systematic variation of population receptive field properties across cortical depth in human visual cortex”. In: NeuroImage 139, pp. 427–438. DOI: 10.1016/j.neuroimage.2016.06.048.

Fracasso, Alessio, Susanne J. van Veluw, Fredy Visser, Peter R. Luijten, Wim Spliet, Jaco J. M. Zwanenburg, Serge O. Dumoulin, and Natalia Petridou (2016). “Lines of Baillarger in vivo and ex vivo: myelin contrast across lamina at 7T MRI and histology”. In: NeuroImage 133, pp. 163–175. DOI: 10.1016/j.neuroimage.2016.02.072.

Fujimoto, Kyoko, Jonathan R. Polimeni, André J. W. van der Kouwe, Martin Reuter, Tobias Kober, Thomas Benner, Bruce Fischl, and Lawrence L. Wald (2014). “Quantitative comparison of cortical surface reconstructions from MP2RAGE and multi-echo MPRAGE data at 3 and 7 Tesla”. In: NeuroImage 90, pp. 60–73. DOI: 10.1016/j.neuroimage.2013.12.012.

Gardner, Justin L. (2010). “Is cortical vasculature functionally organized?” In: NeuroImage 49.3, pp. 1953–1956. DOI: 10.1016/j.neuroimage.2009.07.004.

Glover, Gary H. (2011). “Overview of functional magnetic resonance imaging”. In: Neurosurgery Clinics of North America 22.2, pp. 133–139. DOI: 10.1016/j.nec.2010.11.001.

Goodyear, Bradley G. and Ravi S. Menon (2001). “Brief visual stimulation allows mapping of ocular dominance in visual cortex using fMRI”. In: Human Brain Mapping 14.4, pp. 210–217. DOI: 10.1002/hbm.1053.

Greve, Douglas N. and Bruce Fischl (2009). “Accurate and robust brain image alignment using boundary-based registration”. In: NeuroImage 48.1, pp. 63–72. DOI: 10.1016/j.neuroimage.2009.06.060.

Haenelt, Daniel, Robert Trampel, Shahin Nasr, Jonathan R. Polimeni, Roger B. H. Tootell, Martin I. Sereno, Kerrin J. Pine, Luke J. Edwards, Saskia Helbling, and Nikolaus Weiskopf (2023). “High-resolution quantitative and functional MRI indicate lower myelination of thin and thick stripes in human secondary visual cortex”. In: eLife 12, e78756. DOI: 10.7554/eLife.78756.

Haenelt, Daniel, Nikolaus Weiskopf, Roland Mueller, Shahin Nasr, Jonathan R. Polimeni, Roger B H. Tootell, Martin I. Sereno, and Robert Trampel (2019). “Reliable 3D mapping of ocular dominance columns in humans using GE-EPI fMRI at 7 T”. In: Organization for Human Brain Mapping Annual Meeting (OHBM). Rome, Italy.

Haxby, James V. (2012). “Multivariate pattern analysis of fMRI: the early beginnings”. In: NeuroImage 62.2, pp. 852–855. DOI: 10.1016/j.neuroimage.2012.03.016.

Haynes, John-Dylan and Geraint Rees (2005a). “Predicting the orientation of invisible stimuli from activity in human primary visual cortex”. In: Nature Neuroscience 8.5, pp. 686–691. DOI: 10.1038/nn1445.

Haynes, John-Dylan and Geraint Rees (2005b). “Predicting the stream of consciousness from activity in human visual cortex”. In: Current Biology 15.14, pp. 1301–1307. DOI: 10.1016/j.cub.2005.06.026.

Hollander, Gilles de, Wietske van der Zwaag, Chencan Qian, Peng Zhang, and Tomas Knapen (2021). “Ultra-high field fMRI reveals origins of feedforward and feedback activity within laminae of human ocular dominance columns”. In: NeuroImage 228, p. 117683. DOI: 10.1016/j.neuroimage.2020.117683.

Horton, Jonathan C. and Daniel L. Adams (2005). “The cortical column: a structure without a function”. In: Philosophical Transactions of the Royal Society B: Biological Sciences 360.1456, pp. 837–862. DOI: 10.1098/rstb.2005.1623.

Hubel, David H. and Torsten N. Wiesel (1962). “Receptive fields, binocular interaction and functional architecture in the cat’s visual cortex”. In: The Journal of Physiology 160.1, pp. 106–154. DOI: 10.1113/jphysiol.1962.sp006837.

Hubel, David H. and Torsten N. Wiesel (1969). “Anatomical demonstration of columns in the monkey striate cortex”. In: Nature 221.5182, pp. 747–750. DOI: 10.1038/221747a0.

Huber, Laurentius, Emily S. Finn, Yuhui Chai, Rainer Goebel, Rüdiger Stirnberg, Tony Stöcker, Sean Marrett, Kâmil Uludağ, Seong-Gi Kim, SoHyun Han, Peter A. Bandettini, and Benedikt A. Poser (2021). “Layer-dependent functional connectivity methods”. In: Progress in Neurobiology 207, p. 101835. DOI: 10.1016/j.pneurobio.2020.101835.

Huber, Laurentius, Daniel A. Handwerker, David C. Jangraw, Gang Chen, Andrew Hall, Carsten Stüber, Javier Gonzalez-Castillo, Dimo Ivanov, Sean Marrett, Maria Guidi, Jozien Goense, Benedikt A. Poser, and Peter A. Bandettini (2017). “High-resolution CBV-fMRI allows mapping of laminar activity and connectivity of cortical input and output in human M1”. In: Neuron 96.6, pp. 1253–1263. DOI: 10.1016/j.neuron.2017.11.005.

Huber, Laurentius, Dimo Ivanov, Steffen N. Krieger, Markus N. Streicher, Toralf Mildner, Benedikt A. Poser, Harald E. Möller, and Robert Turner (2014). “Slab-selective, BOLD-corrected VASO at 7 Tesla provides measures of cerebral blood volume reactivity with high signal-to-noise ratio”. In: Magnetic Resonance in Medicine 72.1, pp. 137–148. DOI: 10.1002/mrm.24916.

Huntenburg, Julia M., Christopher J. Steele, and Pierre-Louis Bazin (2018). “Nighres: processing tools for high-resolution neuroimaging”. In: GigaScience 7.7. DOI: 10.1093/gigascience/giy082.

Iamshchinina, Polina, Daniel Kaiser, Renat Yakupov, Daniel Haenelt, Alessandro Sciarra, Hendrik Mattern, Falk Luesebrink, Emrah Duezel, Oliver Speck, Nikolaus Weiskopf, and Radoslaw M. Cichy (2021). “Perceived and mentally rotated contents are differentially represented in cortical depth of V1”. In: Communications Biology 4.1, p. 1069. DOI: 10.1038/s42003-021-02582-4.

Jezzard, Peter and Robert S. Balaban (1995). “Correction for geometric distortion in echo planar images from B0 field variations”. In: Magnetic Resonance in Medicine 34.1, pp. 65–73. DOI: 10.1002/mrm.1910340111.

Julesz, Béla (1971). Foundations of cyclopean perception. Chicago: University of Chicago Press.

Kamitani, Yukiyasu and Frank Tong (2005). “Decoding the visual and subjective contents of the human brain”. In: Nature Neuroscience 8.5, pp. 679–685. DOI: 10.1038/nn1444.

Kay, Kendrick, Keith W. Jamison, Luca Vizioli, Ru-Yuan Zhang, Eshed Margalit, and Kâmil Uğurbil (2019). “A critical assessment of data quality and venous effects in sub-millimeter fMRI”. In: NeuroImage 189, pp. 847–869. DOI: 10.1016/j.neuroimage.2019.02.006.

Kay, Kendrick, Keith W. Jamison, Ru-Yuan Zhang, and Kâmil Uğurbil (2020). “A temporal decomposition method for identifying venous effects in task-based fMRI”. In: Nature Methods 17.10, pp. 1033–1039. DOI: 10.1038/s41592-020-0941-6.

Kennedy, C., M. H. Des Rosiers, O. Sakurada, M. Shinohara, M. Reivich, J. W. Jehle, and L. Sokoloff (1976). “Metabolic mapping of the primary visual system of the monkey by means of the autoradiographic [14C]deoxyglucose technique”. In: Proceedings of the National Academy of Sciences 73.11, pp. 4230–4234. DOI: 10.1073/pnas.73.11.4230.

Kleiner, Mario, David H. Brainard, Denis G. Pelli, Allen Ingling, Richard Murray, and Chris Broussard (2007). “What’s new in psychtoolbox-3”. In: Perception 36.14, pp. 1–16. DOI: 10.1177/03010066070360S101.

Kok, Peter, Lauren J. Bains, Tim van Mourik, David G. Norris, and Floris P. de Lange (2016). “Selective activation of the deep layers of the human primary visual cortex by top-down feedback”. In: Current Biology 26.3, pp. 371–376. DOI: 10.1016/j.cub.2015.12.038.

Koopmans, Peter J., Markus Barth, and David G. Norris (2010). “Layer-specific BOLD activation in human V1”. In: Human Brain Mapping 31.9, pp. 1297–1304. DOI: 10.1002/hbm.20936.

Kriegeskorte, Nikolaus and Peter A. Bandettini (2007). “Analyzing for information, not activation, to exploit high-resolution fMRI”. In: NeuroImage 38.4, pp. 649–662. DOI: 10.1016/j.neuroimage.2007.02.022.

Kriegeskorte, Nikolaus, Rhodri Cusack, and Peter A. Bandettini (2010). “How does an fMRI voxel sample the neuronal activity pattern: compact-kernel or complex spatiotemporal filter?” In: NeuroImage 49.3, pp. 1965–1976. DOI: 10.1016/j.neuroimage.2009.09.059.

LeVay, Simon, David H. Hubel, and Torsten N. Wiesel (1975). “The pattern of ocular dominance columns in macaque visual cortex revealed by a reduced silver stain”. In: Journal of Comparative Neurology 159.4, pp. 559–575. DOI: 10.1002/cne.901590408.

Mandelkow, Hendrik, Jacobus A. Zwart, and Jeff H. Duyn (2017). “Effects of spatial fMRI resolution on the classification of naturalistic movies”. In: NeuroImage 162, pp. 45–55. DOI: 10.1016/j.neuroimage.2017.08.053.

Markuerkiaga, Irati, Markus Barth, and David G. Norris (2016). “A cortical vascular model for examining the specificity of the laminar BOLD signal”. In: NeuroImage 132, pp. 491–498. DOI: 10.1016/j.neuroimage.2016.02.073.

Marquardt, Ingo, Peter De Weerd, Marian Schneider, Omer Faruk Gulban, Dimo Ivanov, Yawen Wang, and Kâmil Uludağ (2020). “Feedback contribution to surface motion perception in the human early visual cortex”. In: eLife 9, e50933. DOI: 10.7554/eLife.50933.

Marques, José P, Tobias Kober, Gunnar Krueger, Wietske van der Zwaag, Pierre-François Van de Moortele, and Rolf Gruetter (2010). “MP2RAGE, a self bias-field corrected sequence for improved segmentation and T1-mapping at high field”. In: NeuroImage 49.2, pp. 1271–1281. DOI: 10.1016/j.neuroimage.2009.10.002.

Menon, Ravi S. and Bradley G. Goodyear (1999). “Submillimeter functional localization in human striate cortex using BOLD contrast at 4 Tesla: implications for the vascular point-spread function”. In: Magnetic Resonance in Medicine 41.2, pp. 230–235. DOI: 10.1002/(SICI)1522-2594(199902)41:2<230::AID-MRM3>3.0.CO;2-O.

Menon, Ravi S., Seiji Ogawa, John P. Strupp, and Kâmil Uğurbil (1997). “Ocular dominance in human V1 demonstrated by functional magnetic resonance imaging”. In: Journal of Neurophysiology 77.5, pp. 2780–2787. DOI: 10.1152/jn.1997.77.5.2780.

Miles, Walter R. (1929). “Ocular dominance demonstrated by unconscious sighting”. In: Journal of Experimental Psychology 12.2, pp. 113–126. DOI: 10.1037/h0075694.

Misaki, Masaya, Wen-Ming Luh, and Peter A. Bandettini (2013). “The effect of spatial smoothing on fMRI decoding of columnar-level organization with linear support vector machine”. In: Journal of Neuroscience Methods 212.2, pp. 355–361. DOI: 10.1016/j.jneumeth.2012.11.004.

Moeller, Steen, Essa Yacoub, Cheryl A. Olman, Edward Auerbach, John Strupp, Noam Harel, and Kâmil Uğurbil (2010). “Multiband multislice GE-EPI at 7 Tesla, with 16-fold acceleration using partial parallel imaging with application to high spatial and temporal whole-brain fMRI”. In: Magnetic Resonance in Medicine 63.5, pp. 1144–1153. DOI: 10.1002/mrm.22361.

Moerel, Michelle, Federico De Martino, Valentin G. Kemper, Sebastian Schmitter, An T. Vu, Kâmil Uğurbil, Elia Formisano, and Essa Yacoub (2018). “Sensitivity and specificity considerations for fMRI encoding, decoding, and mapping of auditory cortex at ultra-high field”. In: NeuroImage 164, pp. 18–31. DOI: 10.1016/j.neuroimage.2017.03.063.

Mountcastle, Vernon B. (1957). “Modality and topographic properties of single neurons of cat’s somatic sensory cortex”. In: Journal of Neurophysiology 20.4, pp. 408–434. DOI: 10.1152/jn.1957.20.4.408.

Mountcastle, Vernon B. (1997). “The columnar organization of the neocortex”. In: Brain 120.4, pp. 701–722. DOI: 10.1093/brain/120.4.701.

Movahedian Attar, Fakhereh, Evgeniya Kirilina, Daniel Haenelt, Kerrin J. Pine, Robert Trampel, LukeJ. Edwards, and Nikolaus Weiskopf (2020). “Mapping short association fibers in the early cortical visual processing stream using in vivo diffusion tractography”. In: Cerebral Cortex 30.8, pp. 4496–4514. DOI: 10.1093/cercor/bhaa049.

Muckli, Lars, Federico De Martino, Luca Vizioli, Lucy S. Petro, Fraser W. Smith, Kâmil Uğurbil, Rainer Goebel, and Essa Yacoub (2015). “Contextual feedback to superficial layers of V1”. In: Current Biology 25.20, pp. 2690–2695. DOI: 10.1016/j.cub.2015.08.057.

Nasr, Shahin, Cristen LaPierre, Christopher E. Vaughn, Thomas Witzel, Jason P. Stockmann, and Jonathan R. Polimeni (2020). “In vivo functional localization of the temporal monocular crescent representation in human primary visual cortex”. In: NeuroImage 209, p. 116516. DOI: 10.1016/j.neuroimage.2020.116516.

Nasr, Shahin, Jonathan R. Polimeni, and Roger B. H. Tootell (2016). “Interdigitated color-and disparity-selective columns within human visual cortical areas V2 and V3”. In: The Journal of Neuroscience 36.6, pp. 1841–1857. DOI: 10.1523/JNEUROSCI.3518-15.2016.

Nasr, Shahin, Jan Skerswetat, Eric D. Gaier, Sarala N. Malladi, Bryan Kennedy, Roger B. H. Tootell, Peter Bex, and David G. Hunter (2025). “Differential impacts of strabismic and anisometropic amblyopia on the mesoscale functional organization of the human visual cortex”. In: The Journal of Neuroscience 45.6, e0745242024. DOI: 10.1523/JNEUROSCI.0745-24.2024.

Nieuwenhuys, Rudolf, Jan Voogd, and Christiaan van Huijzen (2008). The human central nervous system: a synopsis and atlas. Berlin, Heidelberg, Germany: Springer. ISBN: 9783540346845.

Obermayer, Klaus and Gary G. Blasdel (1993). “Geometry of orientation and ocular dominance columns in monkey striate cortex”. In: Journal of Neuroscience 13.10, pp. 4114–4129. DOI: 10.1523/JNEUROSCI.13-10-04114.1993.

Oga, Tomofumi, Tsuguhisa Okamoto, and Ichiro Fujita (2016). “Basal dendrites of layer-III pyramidal neurons do not scale with changes in cortical magnification factor in macaque primary visual cortex”. In: Frontiers in Neural Circuits 10. DOI: 10.3389/fncir.2016.00074.

Palomero-Gallagher, Nicola and Karl Zilles (2019). “Cortical layers: cyto-, myelo-, receptor- and synaptic architecture in human cortical areas”. In: NeuroImage 197, pp. 716–741. DOI: 10.1016/j.neuroimage.2017.08.035.

Parkes, Laura M., Jens V. Schwarzbach, Annemieke A. Bouts, Roel H. R. Deckers, Pim Pullens, Christian M. Kerskens, and David G. Norris (2005). “Quantifying the spatial resolution of the gradient echo and spin echo BOLD response at 3 Tesla”. In: Magnetic Resonance in Medicine 54.6, pp. 1465–1472. DOI: 10.1002/mrm.20712.

Pedregosa, Fabian, Gaël Varoquaux, Alexandre Gramfort, Vincent Michel, Bertrand Thirion, Olivier Grisel, Mathieu Blondel, Peter Prettenhofer, Ron Weiss, Vincent Dubourg, Jake Vanderplas, Alexandre Passos, David Cournapeau, Matthieu Brucher, Matthieu Perrot, and Edouard Duchesnay (2011). “Scikit-learn: machine learning in Python”. In: Journal of Machine Learning Research 12, pp. 2825–2830.

Pelli, Denis G. (1997). “The VideoToolbox software for visual psychophysics: transforming numbers into movies”. In: Spatial Vision 10.4, pp. 437–442. DOI: 10.1163/156856897X00366.

Pfaffenrot, Viktor, Maximilian N. Voelker, Sriranga Kashyap, and Peter J. Koopmans (2021). “Laminar fMRI using T2-prepared multi-echo FLASH”. In: NeuroImage 236, p. 118163. DOI: 10.1016/j.neuroimage.2021.118163.

Poggio, Gian F. (1995). “Mechanisms of stereopsis in monkey visual cortex”. In: Cerebral Cortex 5.3, pp. 193–204. DOI: 10.1093/cercor/5.3.193.

Polimeni, Jonathan R., Bruce Fischl, Douglas N. Greve, and Lawrence L. Wald (2010a). “Laminar analysis of 7T BOLD using an imposed spatial activation pattern in human V1”. In: NeuroImage 52.4, pp. 1334–1346. DOI: 10.1016/j.neuroimage.2010.05.005.

Polimeni, Jonathan R., Douglas N. Greve, Bruce Fischl, and Lawrence L. Wald (2010b). “Depth-resolved laminar analysis of resting-state fluctuation amplitude in high-resolution 7T fMRI”. In: Proceedings International Society for Magnetic Resonance in Medicine. Stockholm, Sweden.

Poser, Benedikt A., Peter J. Koopmans, Thomas Witzel, Lawrence L. Wald, and Markus Barth (2010). “Three dimensional echo-planar imaging at 7 Tesla”. In: NeuroImage 51.1, pp. 261–266. DOI: 10.1016/j.neuroimage.2010.01.108.

Schmidt, Marianna E., Daniel Haenelt, Evgeniya Kirilina, and Nikolaus Weiskopf (2024). “Enhanced specificity for single eye ocular dominance column (ODC) mapping using neurophysiologically informed spatial filtering in ultra-high-resolution fMRI”. In: Society for Neuroscience (SfN) Annual Meeting. Chicago, IL USA.

Sengupta, Ayan, Renat Yakupov, Oliver Speck, Stefan Pollmann, and Michael Hanke (2017). “The effect of acquisition resolution on orientation decoding from V1 BOLD fMRI at 7 T”. In: NeuroImage 148, pp. 64–76. DOI: 10.1016/j.neuroimage.2016.12.040.

Sereno, Martin I., Anders M. Dale, John B. Reppas, Kenneth K. Kwong, John W. Belliveau, Thomas J. Brady, Bruce R. Rosen, and Roger B. H. Tootell (1995). “Borders of multiple visual areas in humans revealed by functional magnetic resonance imaging”. In: Science 268.5212, pp. 889–893. DOI: 10.1126/science.7754376.

Shmuel, Amir, Denis Chaimow, Guenter Raddatz, Kâmil Uğurbil, and Essa Yacoub (2010). “Mechanisms underlying decoding at 7 T: ocular dominance columns, broad structures, and macroscopic blood vessels in V1 convey information on the stimulated eye”. In: NeuroImage 49.3, pp. 1957–1964. DOI: 10.1016/j.neuroimage.2009.08.040.

Shmuel, Amir, Essa Yacoub, Denis Chaimow, Nikos K. Logothetis, and Kâmil Uğurbil (2007). “Spatio-temporal point-spread function of fMRI signal in human gray matter at 7 Tesla”. In: NeuroImage 35.2, pp. 539–552. DOI: 10.1016/j.neuroimage.2006.12.030.

Silva, Afonso C., Alan P. Koretsky, and Jeff H. Duyn (2007). “Functional MRI impulse response for BOLD and CBV contrast in rat somatosensory cortex”. In: Magnetic Resonance in Medicine 57.6, pp. 1110–1118. DOI: 10.1002/mrm.21246.

Stüber, Carsten, Markus Morawski, Andreas Schäfer, Christian Labadie, Miriam Wähnert, Christoph Leuze, Markus Streicher, Nirav Barapatre, Katja Reimann, Stefan Geyer, Daniel Spemann, and Robert Turner (2014). “Myelin and iron concentration in the human brain: a quantitative study of MRI contrast”. In: NeuroImage 93.Pt 1, pp. 95–106. DOI: 10.1016/j.neuroimage.2014.02.026.

Swisher, Jascha D., J. Christopher Gatenby, John C. Gore, Benjamin A. Wolfe, Chan-Hong Moon, Seong-Gi Kim, and Frank Tong (2010). “Multiscale pattern analysis of orientation-selective activity in the primary visual cortex”. In: Journal of Neuroscience 30.1, pp. 325–330. DOI: 10.1523/JNEUROSCI.4811-09.2010.

Talagala, S. Lalith, Joelle E. Sarlls, Siyuan Liu, and Souheil J. Inati (2016). “Improvement of temporal signal-to-noise ratio of GRAPPA accelerated echo planar imaging using a FLASH based calibration scan”. In: Magnetic Resonance in Medicine 75.6, pp. 2362–2371. DOI: 10.1002/mrm.25846.

Tootell, Roger B. H., Nouchine K. Hadjikhani, Wim Vanduffel, Arthur K. Liu, Janine D. Mendola, Martin I. Sereno, and Anders M. Dale (1998). “Functional analysis of primary visual cortex (V1) in humans”. In: Proceedings of the National Academy of Sciences 95.3, pp. 811–817. DOI: 10.1073/pnas.95.3.811.

Tootell, Roger B. H., Susan L. Hamilton, Martin S. Silverman, and Eugene Switkes (1988). “Functional anatomy of macaque striate cortex. I. Ocular dominance, binocular interactions, and baseline conditions”. In: Journal of Neuroscience 8.5, pp. 1500–1530. DOI: 10.1523/JNEUROSCI.08-05-01500.1988.

Tootell, Roger B. H. and Shahin Nasr (2017). “Columnar segregation of magnocellular and parvocellular streams in human extrastriate cortex”. In: The Journal of Neuroscience 37.33, pp. 8014–8032. DOI: 10.1523/JNEUROSCI.0690-17.2017.

Trampel, Robert, Pierre-Louis Bazin, Kerrin J. Pine, and Nikolaus Weiskopf (2019). “In-vivo magnetic resonance imaging (MRI) of laminae in the human cortex”. In: NeuroImage 197, pp. 707–715. DOI: 10.1016/j.neuroimage.2017.09.037.

Trampel, Robert, Derek V. M. Ott, and Robert Turner (2011). “Do the congenitally blind have a stria of Gennari? First intracortical insights in vivo”. In: Cerebral Cortex 21.9, pp. 2075–2081. DOI: 10.1093/cercor/bhq282.

Turner, Robert (2002). “How much cortex can a vein drain? Downstream dilution of activation-related cerebral blood oxygenation changes”. In: NeuroImage 16.4, pp. 1062–1067. DOI: 10.1006/nimg.2002.1082.

Turner, Robert and Stefan Geyer (2014). “Comparing like with like: the power of knowing where you are”. In: Brain Connectivity 4.7, pp. 547–557. DOI: 10.1089/brain.2014.0261.

Tustison, Nicholas J., Brian B. Avants, Philip A. Cook, Yuanjie Zheng, Alexander Egan, Paul A. Yushkevich, and James C. Gee (2010). “N4ITK: improved N3 bias correction”. In: IEEE Transactions on Medical Imaging 29.6, pp. 1310–1320. DOI: 10.1109/TMI.2010.2046908.

Vizioli, Luca, Federico De Martino, Lucy S. Petro, Daniel Kersten, Kâmil Uğurbil, Essa Yacoub, and Lars Muckli (2020). “Multivoxel pattern of blood oxygen level dependent activity can be sensitive to stimulus specific fine scale responses”. In: Scientific Reports 10.1, p. 7565. DOI: 10.1038/s41598-020-64044-x.

Vogt, Cécile and Oskar Vogt (1919). Allgemeine Ergebnisse unserer Hirnforschung. Leipzig, Germany: J.A. Barth.

Waehnert, Miriam, Juliane Dinse, Marcel Weiss, Markus Streicher, Waehnert P., Stefan Geyer, Robert Turner, and Pierre-Louis Bazin (2014). “Anatomically motivated modeling of cortical laminae”. In: NeuroImage 93.2, pp. 210–220. DOI: 10.1016/j.neuroimage.2013.03.078.

Wandell, Brian A. (1995). Foundations of vision. Sinauer Associates.

Weber, Bruno, Anna Lena Keller, Johannes Reichold, and Nikos K. Logothetis (2008). “The microvascular system of the striate and extrastriate visual cortex of the macaque”. In: Cerebral Cortex 18.10, pp. 2318–2330. DOI: 10.1093/cercor/bhm259.

Weiskopf, Nikolaus, Luke J. Edwards, Gunther Helms, Siawoosh Mohammadi, and Evgeniya Kirilina (2021). “Quantitative magnetic resonance imaging of brain anatomy and in-vivo histology”. In: Nature Reviews Physics 3, pp. 570–588. DOI: 10.1038/s42254-021-00326-1.

Yacoub, Essa, Amir Shmuel, Nikos Logothetis, and Kâmil Uğurbil (2007). “Robust detection of ocular dominance columns in humans using Hahn spin echo BOLD functional MRI at 7 Tesla”. In: NeuroImage 37.4, pp. 1161–1177. DOI: 10.1016/j.neuroimage.2007.05.020.

Yang, Jiajia, Laurentius Huber, Yinghua Yu, and Peter A. Bandettini (2021). “Linking cortical circuit models to human cognition with laminar fMRI”. In: Neuroscience & Biobehavioral Reviews 128, pp. 467–478. DOI: 10.1016/j.neubiorev.2021.07.005.

Zaretskaya, Natalia, Jonas Bause, Jonathan R. Polimeni, Pablo R. Grassi, Klaus Scheffler, and Andreas Bartels (2020). “Eye-selective fMRI activity in human primary visual cortex: comparison between 3 T and 9.4 T, and effects across cortical depth”. In: NeuroImage 220, p. 117078. DOI: 10.1016/j.neuroimage.2020.117078.

Zaretskaya, Natalia, Bruce Fischl, Martin Reuter, Ville Renvall, and Jonathan R. Polimeni (2018). “Advantages of cortical surface reconstruction using submillimeter 7 T MEMPRAGE”. In: NeuroImage 165, pp. 11–26. DOI: 10.1016/j.neuroimage.2017.09.060.

